# A highly resolved network reveals the role of terrestrial herbivory in structuring aboveground food webs

**DOI:** 10.1101/2023.10.05.560934

**Authors:** Kayla R. S. Hale, John David Curlis, Giorgia G. Auteri, Sasha Bishop, Rowan L. K. French, Lance E. Jones, Kirby L. Mills, Brian G. Scholtens, Meagan Simons, Cody Thompson, Jordon Tourville, Fernanda S. Valdovinos

**Affiliations:** Department of Ecology & Evolutionary Biology, University of Michigan, Ann Arbor, MI, USA; Department of Integrative Biology, University of Guelph, Guelph, ON, CA; Department of Biology, Missouri State University, Springfield, MO, USA; Department of Ecology & Evolutionary Biology, University of Toronto, Toronto, ON, CA; Department of Plant Biology, University of Illinois at Urbana-Champaign, Urbana, IL, USA; School for Environment & Sustainability, University of Michigan, Ann Arbor, MI, USA; Department of Biology, College of Charleston, Charleston, SC, USA; Museum of Zoology, University of Michigan, Ann Arbor, MI, USA; Department of Environmental Biology, SUNY College of Environmental Science and Forestry, Syracuse, NY, USA; Research Department, Appalachian Mountain Club, Boston, MA, USA; Department of Environmental Science and Policy, University of California, Davis, CA, USA

**Keywords:** Multiplex ecological network, plant-insect interactions, temperate forest ecosystem, niche model, trophic species, scale-dependence

## Abstract

Comparative studies suggest remarkable similarities among food webs across habitats, including systematic changes in their structure with diversity and complexity (scale-dependence). However, historic aboveground terrestrial food webs (ATFWs) have coarsely grouped plants and insects such that these webs are generally small, and herbivory is disproportionately underrepresented compared to vertebrate predator-prey interactions. Furthermore, terrestrial herbivory is thought to be structured by unique processes compared to size-structured feeding in other systems. Here, we present the richest ATFW to date, including ∼580,000 feeding links among ∼3,800 taxonomic species, sourced from ∼27,000 expert-vetted interaction records annotated as feeding upon one of six different resource types: leaves, flowers, seeds, wood, prey, and carrion. By comparison to historical ATFWs and null ecological hypotheses, we show that our temperate forest web displays a potentially unique structure characterized by two properties: a) a large fraction of carnivory interactions dominated by a small number of hyper-generalist, opportunistic bird and bat predators, and b) a smaller fraction of herbivory interactions dominated by a hyper-rich community of insects with variably-sized but highly-specific diets. We attribute our findings to the large-scale, even resolution of vertebrate, insect, and plant guilds in our food web.

## 1) Introduction

Ecosystems contain immense biological complexity. Food webs represent part of this complexity by documenting the feeding interactions (links) between taxa (nodes). Comparative studies of food webs across habitats have revealed robust and nonrandom patterns suggestive of an underlying architecture of life [1–7]. For example, the structure of food webs changes systematically with diversity and complexity (termed “scale-dependence”) but maintains hierarchy, demonstrated in aquatic and belowground food webs through modular, size-structured pathways of larger consumers feeding on smaller resources [8–13]. Aboveground terrestrial food webs (ATFWs) also exhibit size-structure in predator-prey interactions, but different mechanisms (e.g., chemical composition, trait matching) likely underlie the specialized feeding of insect herbivores on often-larger terrestrial plant resources [14]. However, there are few published ATFWs and even fewer that include high-resolution data for both plant-herbivore and predator-prey interactions across broad taxonomic groups (Table 1). Therefore, whether and how the structure of ATFWs may fundamentally differ from those of other habitats remains unclear [15]. As a step towards answering this question, we construct the most extensive ATFW to date and study the mechanisms by which its increased taxonomic and trophic resolution lead to a unique structure relative to the scale-dependent pattern observed in previous webs.

**Table 1.**
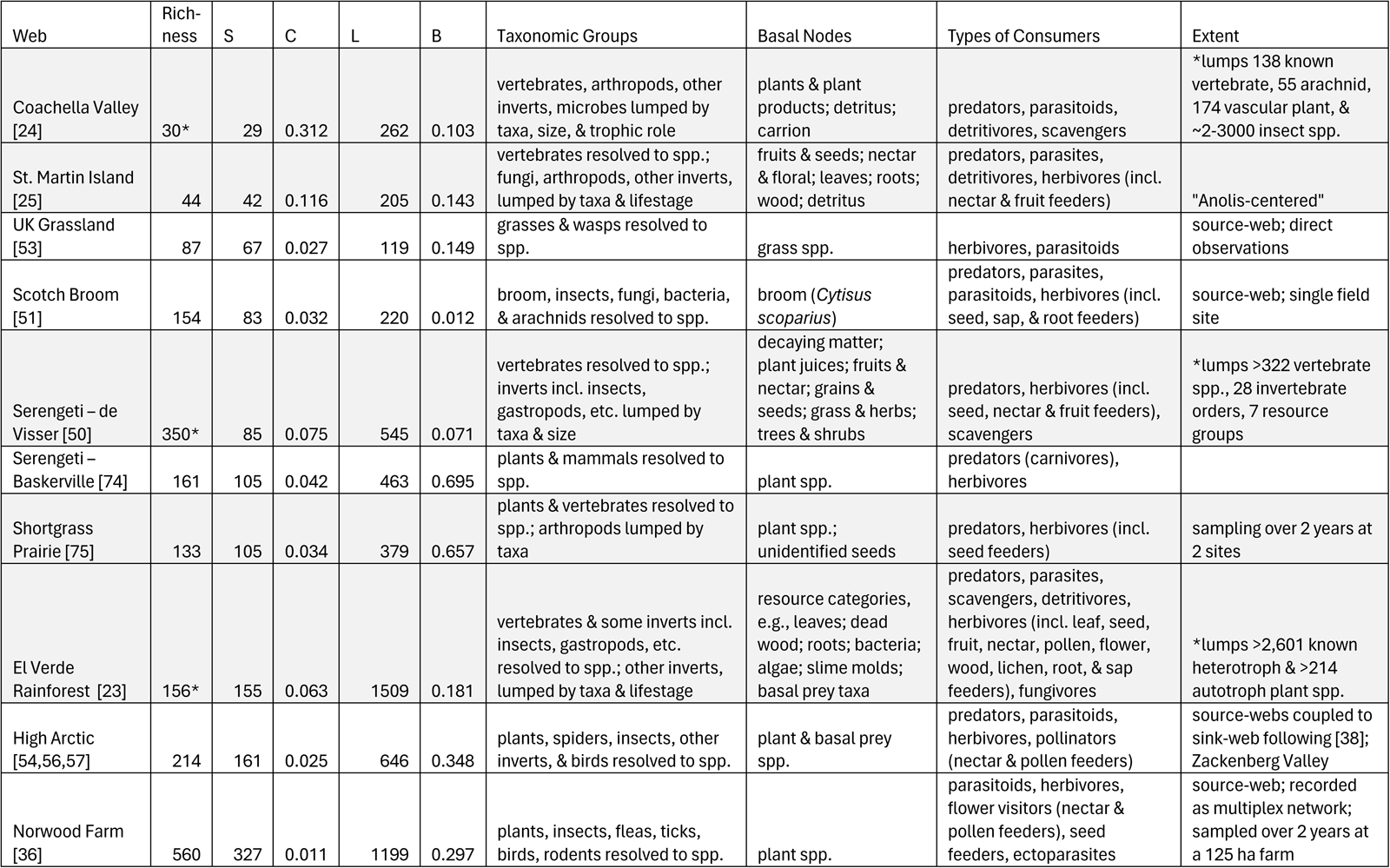
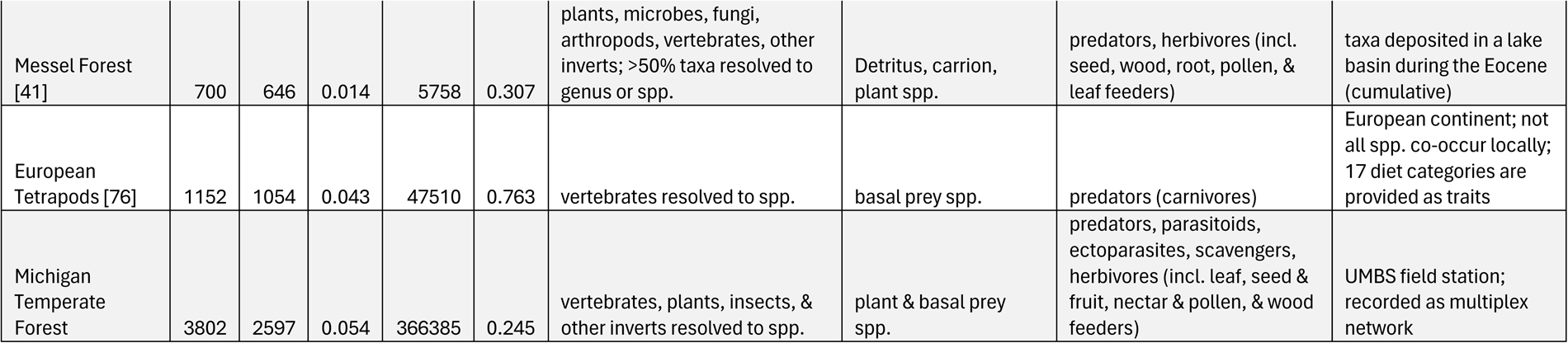
Properties of aboveground terrestrial food webs. A collection of classic and more recent food webs used for comparison to our new Michigan Temperate Forest web for the University of Michigan Biological Station. We include the classic webs (published prior to 2000) traditionally considered ‘highly resolved;’ this excludes the many historical webs in the ECoWeB catalogue of low richness and variable resolution [27]. We exclude modern webs in “container habitats” (e.g., under a log, in a tree hole) as well as newer parasitoid-host webs [6,73]. The High Arctic web was assembled from data in Appendix S2 of Wirta et al. [54], originally from Roslin et al. and Rasmussen et al. [56,57]. Column definitions: *Web:* Traditional name and reference. *Richness*: Number of ‘species’ in original publication (but see *Extent*). *S*: Number of trophic species. *C:* Directed connectance. *L:* Number of links. *B:* Fraction of basal trophic species (with no consumers). *Taxonomic Groups*: Taxa and resolution in the food web. *Basal Nodes:* Types of trophic species or functional groups at the base of the food web. *Types of Consumers:* Types of feeding interactions included in the food web. *Extent:* Notes on the space and time of food web construction. Unless otherwise noted, webs are *cumulative metawebs,* built from records pooled across time (including published literature) and similar habitats (usually contiguous field sites). Other definitions: “lumped by taxa”: grouped to order or family except potentially for key species. *Source-web:* web recording the food chain(s) up from a set of resources. *Sink-web*: web recording the food chain(s) down from a set of consumers. “*Breadth”* webs (highlighted gray) are those that include multiple taxonomic groups and energy pathways but with lower resolution. “*Depth”* webs are those with higher resolution but missing key structural components, including some non-traditional food webs like the Pocock Farm multiplex network. Abbreviations: “incl.” including, “inverts” invertebrates. See text for other definitions.

| Web | Rich-<br>ness | S | C | L | B | Taxonomic Groups | Basal Nodes | Types of Consumers | Extent |
| --- | --- | --- | --- | --- | --- | --- | --- | --- | --- |
| Coachella Valley [24] | 30* | 29 | 0.312 | 262 | 0.103 | vertebrates, arthropods, other inverts, microbes lumped by taxa, size, & trophic role | plants & plant products; detritus; carrion | predators, parasitoids, detritivores, scavengers | *lumps 138 known vertebrate, 55 arachnid, 174 vascular plant, & ~2-3000 insect spp. |
| St. Martin Island [25] | 44 | 42 | 0.116 | 205 | 0.143 | vertebrates resolved to spp.; fungi, arthropods, other inverts, lumped by taxa & lifestage | fruits & seeds; nectar & floral; leaves; roots; wood; detritus | predators, parasites, detritivores, herbivores (incl. nectar & fruit feeders) | "Anolis-centered" |
| UK Grassland [53] | 87 | 67 | 0.027 | 119 | 0.149 | grasses & wasps resolved to spp. | grass spp. | herbivores, parasitoids | source-web; direct observations |
| Scotch Broom [51] | 154 | 83 | 0.032 | 220 | 0.012 | broom, insects, fungi, bacteria, & arachnids resolved to spp. | broom (*Cytisus scoparius*) | predators, parasites, parasitoids, herbivores (incl. seed, sap, & root feeders) | source-web; single field site |
| Serengeti – de Visser [50] | 350* | 85 | 0.075 | 545 | 0.071 | vertebrates resolved to spp.; inverts incl. insects, gastropods, etc. lumped by taxa & size | decaying matter; plant juices; fruits & nectar; grains & seeds; grass & herbs; trees & shrubs | predators, herbivores (incl. seed, nectar & fruit feeders), scavengers | *lumps >322 vertebrate spp., 28 invertebrate orders, 7 resource groups |
| Serengeti – Baskerville [74] | 161 | 105 | 0.042 | 463 | 0.695 | plants & mammals resolved to spp. | plant spp. | predators (carnivores), herbivores |  |
| Shortgrass Prairie [75] | 133 | 105 | 0.034 | 379 | 0.657 | plants & vertebrates resolved to spp.; arthropods lumped by taxa | plant spp.; unidentified seeds | predators, herbivores (incl. seed feeders) | sampling over 2 years at 2 sites |
| El Verde Rainforest [23] | 156* | 155 | 0.063 | 1509 | 0.181 | vertebrates & some inverts incl. insects, gastropods, etc. resolved to spp.; other inverts, lumped by taxa & lifestage | resource categories, e.g., leaves; dead wood; roots; bacteria; algae; slime molds; basal prey taxa | predators, parasites, scavengers, detritivores, herbivores (incl. leaf, seed, fruit, nectar, pollen, flower, wood, lichen, root, & sap feeders), fungivores | *lumps >2,601 known heterotroph & >214 autotroph plant spp. |
| High Arctic [54,56,57] | 214 | 161 | 0.025 | 646 | 0.348 | plants, spiders, insects, other inverts, & birds resolved to spp. | plant & basal prey spp. | predators, parasitoids, herbivores, pollinators (nectar & pollen feeders) | source-webs coupled to sink-web following [38]; Zackenberg Valley |
| Norwood Farm [36] | 560 | 327 | 0.011 | 1199 | 0.297 | plants, insects, fleas, ticks, birds, rodents resolved to spp. | plant spp. | parasitoids, herbivores, flower visitors (nectar & pollen feeders), seed feeders, ectoparasites | source-web; recorded as multiplex network; sampled over 2 years at a 125 ha farm |
| Messel Forest [41] | 700 | 646 | 0.014 | 5758 | 0.307 | plants, microbes, fungi, arthropods, vertebrates, other inverts; >50% taxa resolved to genus or spp. | Detritus, carrion, plant spp. | predators, herbivores (incl. seed, wood, root, pollen, & leaf feeders) | taxa deposited in a lake basin during the Eocene (cumulative) |
| European Tetrapods [76] | 1152 | 1054 | 0.043 | 47510 | 0.763 | vertebrates resolved to spp. | basal prey spp. | predators (carnivores) | European continent; not all spp. co-occur locally; 17 diet categories are provided as traits |
| Michigan Temperate Forest | 3802 | 2597 | 0.054 | 366385 | 0.245 | vertebrates, plants, insects, & other inverts resolved to spp. | plant & basal prey spp. | predators, parasitoids, ectoparasites, scavengers, herbivores (incl. leaf, seed & fruit, nectar & pollen, & wood feeders) | UMBS field station; recorded as multiplex network |

Constructing food webs is fraught with methodological difficulties [16,17]. Observations of species and interactions depend on the boundaries of the system, the specific spatial (vertical versus horizontal transects, microhabitats) and temporal (seasonal, diurnal, duration) scales of sampling, as well as the taxonomic expertise of the investigators (including ability to detect and identify both consumer and resource species). Many organisms regularly cross ecosystem boundaries as part of their life cycles; for example, some insect species spend larval stages underground or underwater, then move to aboveground habitats after maturation, after which they may migrate to a completely different region for breeding [18]. Species also exhibit adaptive foraging and defensive behaviors, effectively “rewiring” interactions in response to changing biotic and abiotic conditions [14,19,20]. One approach to these problems is to construct an expert-vetted “cumulative” or “meta” food web that pools all species and interactions recorded across time and/or similar habitats [21]. This reduces the likelihood of missing cryptic or rare species and provides a more comprehensive accounting of all potential feeding interactions in the system. Additionally, as human activities alter species’ distributions and habitats, “rare” and novel interactions are increasing in frequency [19], making cumulative webs even more important.

Even cumulative food webs rely heavily on expertise and long-term and/or regional sampling. Perhaps for this reason, previous high-quality ATFWs have tended to focus either on taxonomic *breadth* or *depth*. Webs with taxonomic *breadth* (Table 1, gray highlighted rows) tend to resolve vertebrates most highly, while aggregating invertebrates and plants into coarse taxonomic or functional groups (e.g., into insect orders or plant tissue categories). This ‘lumping’ strategy *sensu* Briand [22] seeks to describe broad system-level behavior but is largely a result of the technical difficulties associated with documenting and representing the pure volume of plant-insect associations [23]. Nevertheless, classic breadth webs – Coachella Valley [24], St. Martin Island [25], and El Verde Rainforest [23] – have proven highly influential and remain perhaps our best description of ATFWs because they used cumulative approaches with known species lists from long-term fields sites. Indeed, these webs, with the Little Rock Lake web of Martinez [26], contributed to overturning ‘empirical generalizations’ (such as scale-invariance, low omnivory, etc.) derived from a catalogue of less resolved webs [27–29].

In contrast, webs with greater *depth* of resolution tend to have narrower scope (Table 1). These webs generally focus only on a single taxonomic group (e.g., only tetrapods) or on a single energetic pathway (e.g., ‘source’ or ‘sink’ webs) [21]. In the same vein, the explosion of ecological networks research in the last two decades has tended to focus on highly specific single interaction types such as frugivory or scavenging, demonstrating the unique structure and importance of these subnetworks for ecosystem dynamics and function [30–33]. However, different subnetworks are rarely recorded in the same system, and as such, it is unknown how they may connect with each other or to their broader food web [30]. An exception is the few ‘multiplex’ networks that report high resolution interactions of different types (i.e., feeding on different resources or with different interaction outcomes) among non-disjoint sets of species [34,35]. These studies bring together interactions that otherwise rarely co-occur in food webs [31–33], especially mutualisms, such as pollination or seed dispersal, which can have a feeding component via consumption of nectar and pollen or seeds and fruits, with other forms of ‘antagonistic’ herbivory such as phloem-feeding by aphids [36–38].

Comparative studies attempt to standardize these diverse approaches to constructing food webs in three ways [21,39]. First, they aggregate empirical webs to “trophic species” webs, where taxa with the same set of consumers and resources are grouped into the same node [28]. This reduces methodological biases within and between webs by retaining only functionally distinct units with unique trophic niches [2]. Second, they compare the properties of trophic species webs to null expectations provided by the well-known “niche model” of Williams and Martinez [8], which embodies specific ecological hypotheses for the mechanisms structuring food webs [40]. This approach provides scale-dependent expectations for food web properties (i.e., given their richness and complexity), and deviations from null expectations (sometimes called “errors”) can be interpreted as rejecting the underlying hypotheses. However, errors are also scale-dependent, meaning that the properties of empirical webs increasingly deviate from niche model expectations with increasing richness [39]. Therefore, third, comparative studies extrapolate from scale-dependent errors to assess whether a focal web exhibits unique properties compared to other webs, given its scale [39,41].

In this study, we leveraged more than a century of research at a biological research station to build the Michigan Temperate Forest food web, the richest ATFW to date. We used a cumulative approach, incorporating public records and occurrence data, supplemented and vetted by experts for local plausibility given species’ traits and behaviors. This resulted in ∼580,000 feeding links among ∼3,800 taxonomic species, represented in a multiplex network according to feeding on different resource types (“prey,” “carrion,” “leaves,” “flowers,” “seeds,” or “wood”). Using comparative food web methods, we (1) characterized the properties of the Michigan Temperate Forest, (2) studied whether the increased taxonomic and trophic resolution leads to unique structure, given its scale, compared to previous ATFWs and, if so, (3) identified potential mechanisms underlying the structure of more or less resolved webs.

## 2) Methods

### a) Site, species list, and feeding records

The University of Michigan Biological Station (UMBS) was established in 1909 on ∼10,000 acres of logged and burned land in northern lower Michigan, USA (45°35.5′ N, 84°43′ W) and represents a strongly seasonal system with historically cold, snowy winters and hot, humid summers. The site has, in recent years, been restored to predominantly dry-mesic, northern hardwood forests with patches of wooded wetlands (hardwood conifer swamp) [42]. A full description of the UMBS site and extended methods are available in the Electronic Supplementary Materials.

Briefly, experts (generally, the authors) vetted and approved species lists of mammals, amphibians, reptiles, vascular plants, birds, insects, and non-insect arthropods from UMBS records. These were accumulated from resident biologists’ personal observations, student projects, museum and herbarium specimens, and semi-regular BioBlitz events, in which teams of biologists roamed the site and identified as many organisms as possible. Hereafter, we refer to all approved taxa as “species,” though a small fraction (4.5%) are genera.

The same experts vetted and annotated a list of potential feeding interactions, sourced from region-specific field guides and online databases [43]. Each focal taxon was resolved to species-level, but their interaction partners could be recorded at any taxonomic level (e.g., species *x* eats family *y*). We included all records of direct interactions among species in our system with a bioenergetic flow (i.e., one species consuming another), regardless of lifestage or potential ecological effects (i.e., whether potentially “mutualistic” or “antagonistic”, see [14]. Experts approved recorded interactions between species as plausible if the species co-occur (with respect to phenology, activity patterns, microhabitat usage) and have no trait incompatibilities (with respect to acquisition, ingestion, and assimilation). If a partner in a potential interaction was recorded at a coarser taxonomic level than species, the record was approved only if these conditions could also plausibly hold for all local species in that taxonomic unit.

Lastly, experts categorized records by their focal resource type as animal tissues, either (1) live tissues and as prey, or (2) scavenged as carrion, carcasses, or other decaying animal remains, or as plant tissues, grouped as (3) leaves and stems, (4) flowers, nectar, pollen, etc., (5) seeds, fruits, etc., or (6) wood and bark. Hereafter, we refer to these resource types simply as “prey,” “carrion,” “leaves,” “flowers,” “seeds,” and “wood,” respectively.

### b) Network representation

To translate our list of feeding records into a food web, we began with a “multiplex network” approach (Fig. 1A) in which feeding on different resource types is represented by different types of links between the same set of nodes (taxonomic species). This allowed us to distinguish between the niches of animals feeding on different resources while also accounting for the fact that such resources are coupled together in the same organism. Specifically, we defined a node for each focal species *i* in our list. Then, we defined a directed link of type *l* between nodes *i*, *j* if *i* consumes tissue type *l* of *j* or tissue type *l* of a broader taxonomic group including *j*. Links are binary, indicating the presence or absence of potential feeding, not its frequency, probability, rate, or strength. We retained only unique links, but tracked the most resolved taxonomic level from which each link was sourced. We characterized the complexity of the multiplex network by counting the number of links (*L_l_*), consumer species (*A_l_*), and resource species (*P_l_*) involved in feeding of type *l*, and the fraction of the maximum possible links that were realized as either bipartite connectance = *L_l_*/(*A_l_P_l_*) for feeding on plant tissues (leaves, flowers, seeds, or wood) or unipartite connectance = *L_l_*/(*A_l_* + *P_l_*)^2^ for feeding on animal tissues (prey or carrion).

**Figure 1.**
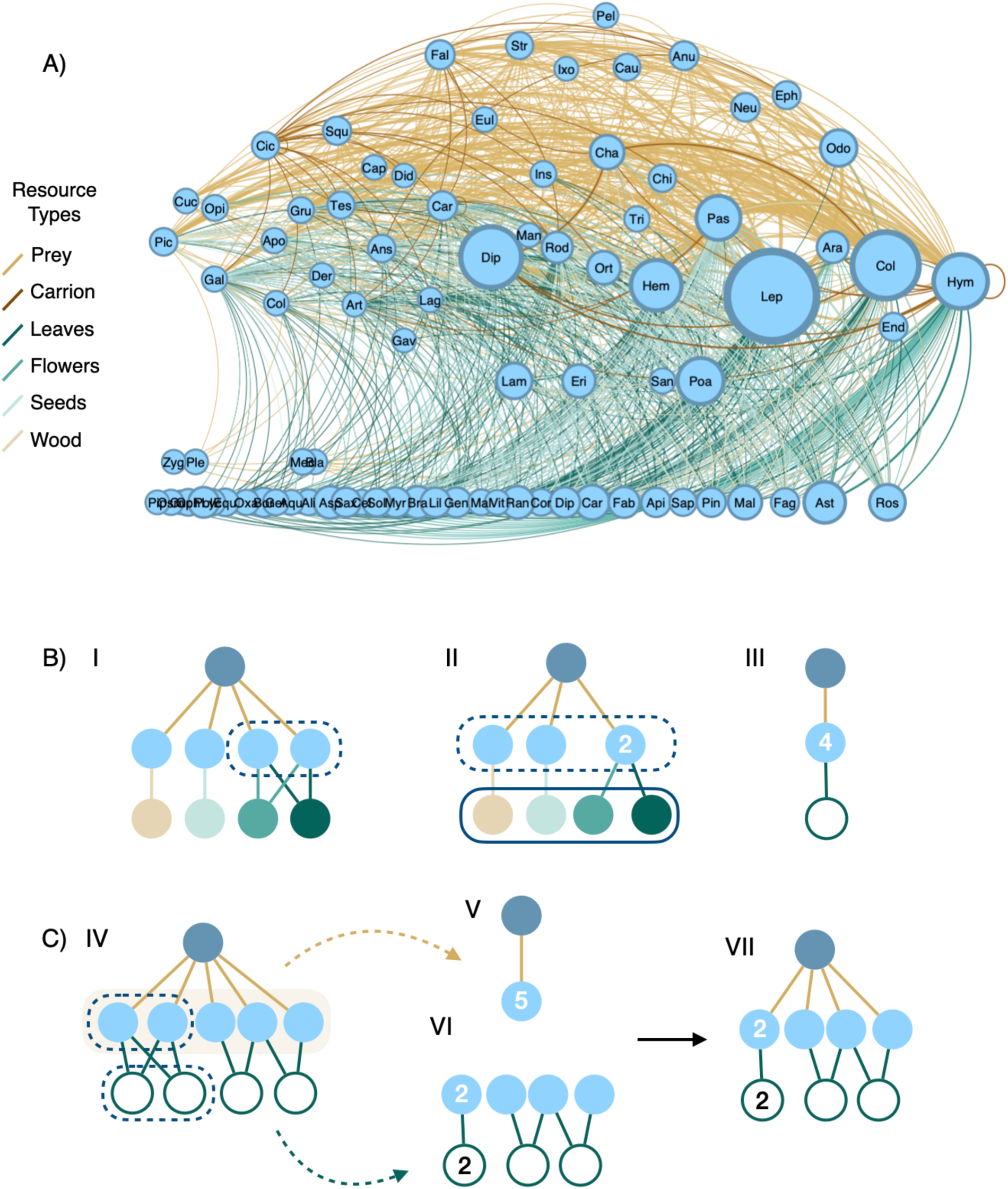
Representations of the Michigan Temperate Forest food web. **(A)** Simplified visualization of the multiplex food web. Each node represents a taxonomic order (labeled by its first three letters), with width scaled to the number of local species, ranging from one (e.g., order Gaviiformes, represented only by the Common Loon) to >1,000 (order Lepidoptera). Each link is a feeding interaction between orders, with width scaled to the total number of feeding interactions between species in each order. Links colored by the resource type consumed as follows: tan – live prey and animal tissues; brown – scavenged carrion; dark green – leaves and stems; green – floral resources; light green – seeds and fruits; and beige – wood and bark. Self-links indicate feeding among species within the order, including cannibalism. Nodes are ordered horizontally by their number of consumers (in-degree), increasing from left to right, and vertically by increasing trophic level (TL) from basal resources on the bottom (TL = 1) to carnivores at the top. Three carnivorous/parasitic plant orders were assigned TL = 1.75 and four basal animal orders were assigned trophic levels 1.25 for visualization. The 85 orders shown here represent 3,082 taxonomic species. **(B)** Illustrated effect of aggregating the multiplex network into ‘trophic species’ – species with same set of consumers and resources. Numbers indicate how many nodes were aggregated into a trophic species. Here, one generalist predator feeds on four herbivores of the same plant species, with its different tissue types (wood, seeds, flowers, and leaves) illustrated as different nodes for clarity (colors follow panel A). There are six taxonomic multiplex species total in this example. The ‘multiplex network’ representation **(I)** can differentiate among the trophic niches of herbivores feeding on different plant tissues. Aggregating into a ‘trophic-species multiplex network’ **(II)** groups the two herbivores that feed on the same plant tissues into one trophic species, resulting in five multiplex trophic species total. Further aggregating all four herbivores into one trophic species feeding on the plant species (white node), results in three trophic species total. This is the traditional representation of ‘trophic-species food webs’ **(III)**, where only taxonomic identity differentiates species’ trophic niches. **(C)** Illustration of the disproportionate effect of herbivory on trophic species resolution in the Michigan Temperate Forest (MTF) web. The generality of predators (especially birds and bats) in the MTF web **(IV)** means that prey species (especially insect herbivores) tend to be aggregated into fewer trophic species when considering carnivory interactions alone **(V)**. The specificity of herbivores in the MTF web results in more trophic species resolved when considering herbivory interactions alone **(VI)**. This leads to a pattern of numerous generalized carnivory interactions and fewer but more specific herbivory interactions among the more numerous plant and herbivore species **(VII)**.

As a direct comparison to previously published food webs (Table 1), we used the conventional food web approach, where a binary link occurs between nodes *i*, *j* if *i* consumes *j* in the multiplex network (i.e., consumes any resource type of species *j*). Following convention, we aggregated all webs into “trophic species” versions, wherein taxa with the same sets of consumer and resource species are grouped together (Fig. 1BIII).

To study the contribution of resolving feeding on different resource types to food web structure, we aggregated our multiplex network to a “trophic species multiplex network” (Fig. 1BI-II), wherein all taxa with the same set of consumers and resources, both in terms of taxonomic species and resource types, are grouped into a single node. The difference in resulting richness between this trophic species multiplex network (Fig. 1BII) and that of our trophic species food web (Fig. 1BIII) indicates how many taxonomic species’ trophic niches are differentiated only by feeding on specific resource types (e.g., on the leaves versus the flowers or the live prey versus the carrion of the same resource species).

Hereafter, we discuss the structure of a “food web” as the trophic-species food web with richness denoted as *S*, number of links denoted as *L*, and directed connectance (the fraction of observed links out of the maximum possible, *L*/*S*^2^) denoted as *C* [26].

### c) Food web structure

To provide null expectations for the scale-dependent structure of food webs, we used the niche model of Williams & Martinez [8] to simulate *N* = 1,000 matching webs using the *S* and *C* of our food web and each of the previous webs (Table 1). Traditionally, the model assumes that all nodes are unique trophic species and webs that include nodes with the same consumer and resource set (i.e., duplicate trophic species) are rejected. For webs with low connectance, we slightly relaxed that assumption and allowed niche model webs to be seeded with a slightly higher initial richness as long as (following another trophic species aggregation) *S* and *C* matched the empirical trophic-species web.

We calculated a suite of properties to characterize the Composition, Hierarchy, and Degree Distribution of the empirical food webs [8,39,44–46]. See Supplementary Table 1 for a full list of properties and definitions. For each structural property, we assessed the significance of deviations from null expectations using normalized model errors (NMEs) [44,47]. NMEs are calculated as the difference between the median model and empirical values normalized by either the difference between the median model value and the 97.5 percentile of the model distribution if the empirical value is greater than the model median, or, if the empirical value is less than the model median, by the difference between the 2.5 percentile of the model distribution and the model median. Values >1 or <–1 indicate that the empirical value is significantly higher or lower, respectively, than the null expectation at the 95% confidence level. For the purposes of discussion, we follow previous works to summarize these as a composite *mean* |NME|, though the individual properties are not independent (see Supplementary Methods) [39].

To characterize Species Composition, we calculated the *fraction of trophic species* in each of the following categories: *Basal (B)*: with consumers but no resources; *Intermediate (I)*: with both consumers and resources; *Top (T)*: with resources but no consumers; *Herbiv (TL2):* eat only basal species (are strict herbivores, i.e., trophic level [TL] = 2); *Carniv:* eat only other consumers (strict carnivores); *Omniv:* eat both basal and consumer species (omnivores); *Cannib:* eat members of their own species (cannibals).

To characterize Link Composition, we calculated *HerbLink,* the fraction of total feeding links that are herbivorous (i.e., are on basal resources), and *TL2Link,* the fraction of feeding links from TL2 herbivores.

To characterize Hierarchy, we calculated *meanTL and maxTL,* the mean and maximum short-weighted trophic level of consumers [46], and *meanTLTop*, the mean trophic level of top consumers.

Degree describes the number of resources (in-degree) and consumers (out-degree) of a species. To characterize Degree Distribution, we calculated *meanGen*, the mean in-degree of consumers (i.e., their “generality”), *GenSD*, the normalized variability of generality, *meanVul*, the mean out-degree of resources (i.e., their “vulnerability”), and *VulSD*, the normalized variability of vulnerability. We also calculated these properties for specific subgroups of species to characterize their respective contribution to deviations observed from null expectations.

Finally, we used two-sample *Kolmogorov-Smirnov* tests to directly test the null hypothesis that the observed and expected degree distributions were sampled from the same underlying distribution [47]. Full results are reported in Supplementary Table 2.

We observed that the niche model tended to generate webs with low *B* at high *S*, which may drive substantial NMEs simply due to underlying differences in species composition. We directly tested whether this may be the case by simulating *N =* 1,000 niche model webs with matching *S*, *C*, *B* values to our empirical food web, called the “basal-matched” treatment. We seeded the niche model with *S* = 2,896, *C* = 0.057, and set the 927 lowest niche-value species to have feeding ranges of zero (thus forcing them to be basal).

All data cleaning and network analyses were performed in MATLAB R2021b [48].

## 3) Results

### a) Species list and feeding records

Our final species list includes 3,802 local species, representing 2,073 genera in 451 families of 85 orders (Fig. 1A). Insects (2,669 spp.) and vascular plants (781 spp.) numerically dominate the community, accounting for ∼90% of the taxa, compared to vertebrates (313 spp.) and non-insect arthropods (39 spp.). The richest orders are insects, especially the Lepidoptera (butterflies and moths, 1,168 spp.), Coleoptera (beetles, 512 spp.), Diptera (flies, 390 spp.), Hymenoptera (bees, wasps, and ants, 265 spp.), and Hemiptera (true bugs, 211 spp.). Worldwide, there are more than twice as many named species of Coleoptera as Lepidoptera [49]; therefore, Lepidopterans are likely substantially overrepresented in our list. Vascular plants represent the most taxonomically diverse group on our list, with 38 orders; however, most of these species belong either to Poales (grasses, sedges, and rushes, 149 spp.) or Asterales (composite flowers, 96 spp.). Vertebrate species include birds (226 spp., including 127 passeriform birds), mammals (52 spp., including 7 bats), amphibians (18 spp.), and reptiles (17 spp.). Finally, non-insect arthropods primarily include spiders and mites, but overall, this group is significantly underrepresented in our list, both in terms of richness and taxonomic diversity.

In sum, we recorded 2,541 species of consumers (including 4 carnivorous or parasitic plants) and 3,782 species of resources. Approximately, two-thirds of species (69.7%) included in the records are resolved to taxonomic species level, while 13.7%, 13.1%, and 3.0% of species (primarily insects) have records from genus-, family-or order-level records at best, respectively. We have no feeding records for 19 plants (0.5% of local species), including most of the Lycopodiales (clubmosses, 5 of 6 spp.) and the Polypodiales (ferns, 11 of 19 spp.), which represent two of the major groups of non-seed plants in our system. Additionally, our records include no diet information for 485 species of insects (16.3% of local animals), primarily from the richest orders (183 Lepidopterans, 145 Dipterans, 109 Coleopterans), but also including all Blattodea (cockroaches, 2 spp.), Plecoptera (stoneflies, 2 spp.), Mecoptera (scorpionflies, 4 spp.), and Zygentoma (silverfish, 1 spp.). Some of these do not feed in aboveground terrestrial habits (or at all) during a certain life stage or feed entirely upon resources we excluded (fungi, detritus, lichens, etc.), limiting their potential diet in our food web. However, these gaps in our dataset may also indicate broader gaps in our expertise or the available natural history information for these species.

We constructed our multiplex network from 26,742 approved records of local taxa feeding on prey, carcass, leaf, flower, seed, or wood resources, totaling 588,416 unique feeding interactions (links) between local species. These links primarily consist of feeding on prey (89.2%), especially insect prey, with the remaining links consisting of scavenging carcasses (0.54%) or feeding on plant leaves (6.7%), flowers (2.2%), seeds (1.2%), or wood (0.17%). Though numerous, these interactions are only a small fraction of the possible links. Separating feeding on each type of resource, we calculate that only 5.8% of the interactions are realized among the 3,023 predator and prey species in our carnivory subnetwork, with similarly low connectances for scavenging (0.2% among 1,357 spp.) and the different types of herbivory (leaves: 3.1% among 2,458 spp., flowers: 2.2% among 1,533 spp., seeds: 5.1% among 796 spp., wood: 9.8% among 200 spp.).

Carnivory is the most numerically dominant interaction in our food web, but only approximately one-third (35.9%) of consumers feed on prey. In fact, the 120 most generalist species in our food web (4.7% of consumers) contribute over half (51.5%) of the unique carnivory links in our network, sourced from only 950 (3.6%) records of focal birds and bats thought to feed opportunistically upon entire insect orders. Records of feeding on insect orders by any taxon contribute 87.1% of unique carnivory links overall, meaning that they are not otherwise included by records at lower taxonomic levels. In comparison, feeding between vertebrates accounts for only 1.3% of carnivory links.

Herbivory interactions are less numerous than carnivory, but most consumers (85.4%) in our food web feed on plants, with nearly half (46.0%) feeding on a single plant tissue. These are primarily insects, dominated by lepidopterans eating leaves (as caterpillars), but also including hymenopterans and dipterans eating floral resources. Over half of herbivory interactions (52.4%) stem from records of feeding between insects and plants at the genus-and species-level, with only 4.1% of unique herbivory interactions contributed by order-level records across all taxonomic groups. Therefore, in contrast to carnivory, our herbivory records at coarser taxonomic levels do not include or are redundant to interactions from more taxonomically-resolved records.

Over one-third of consumers (39.6%) feed upon more than one type of resource. Around half of these feed on leaves and flowers (18.9% of consumers, primarily lepidopterans and coleopterans). A smaller fraction (11.6%) feed on more than two types of resources, but these represent a more diverse set of insects, mammals, and birds feeding on prey, leaves, and flowers or seeds, or, less frequently, leaves, flowers, and wood. Ants (Formicidae, 48 spp.) uniquely feed on prey, leaves, flowers, and carrion. Among consumers feeding on multiple resource types, we observed significant positive correlations between diet breadths when feeding on prey and plant leaves (Pearson correlation: r = 0.26, p = 2.7 x 10^-6^, N = 319), prey and seeds (r = 0.14, p = 0.044, N = 202), leaves and seeds (r = 0.42, p = 7.2 x 10^-9^, N = 178), and leaves and wood (r = 0.24, p = 0.017, N = 98; Supplementary Fig. 1). In other words, generalists on one resource type also tend to be generalists on others. In contrast to the tissue specialization by most animals, most plants (91.3% of 781 species) support consumers on more than one of their tissues, with a small set of diverse plants (81 species in 13 orders) sustaining feeding on all four recorded tissue types (Supplementary Fig. 2).

### b) Trophic species composition

Our final food web consists of *S* = 2,595 trophic species and *L =* 365,951 links. An additional 29 trophic species and 8,462 links would be distinguished by feeding only on different types of plant tissues (Fig. 1B). Nearly all (92.3%, 2,394) of the trophic species nodes correspond to taxonomic species, including all vertebrates, nearly all non-insect arthropods (94.9%), over two-thirds of plants (70.2%), and over half of insects (55.0%) in our species list. The remaining trophic species are each composed of taxonomic species from a single order. This is likely because entire insect orders, the primary resources in the food web, share the same opportunistic/generalist predators and are therefore only distinguished by their diets (Fig. 1C). Over half of these taxonomic species (54.1%, 761) are represented in only 27 trophic species groups; these comprise most (83.1%) of the 485 animal species without diet information (that is, the species most likely lacking resolution).

Feeding on leaves contributed to distinguishing the most trophic species in our food web (924 trophic species), followed by feeding on prey, flowers, seeds, wood, and then carrion (417, 322, 13, 5, and 0 respective additional trophic species relative to versions of the food web built with feeding only on the other resource types). Therefore, herbivory interactions, and particularly leaf herbivory, provide the key component for distinguishing species’ trophic niches in our system.

### c) Scale-dependence of food web structure

The structure of our food web deviates significantly from the null expectation provided by the niche model in almost every metric. Beginning with trophic species composition (Fig. 3A-E), approximately one-quarter of trophic species in our web are basal (*B_obs_* = 0.245) and the remaining three-quarters are intermediate (*I_obs_* = 0.755). Nearly half of trophic species are herbivores that eat only basal trophic species (i.e., are trophic level 2, *TL2_obs_* = 0.483), and nearly one-quarter are omnivores that eat both basal and consumer trophic species (*Omniv_obs_* = 0.235), while <5% are carnivores that feed strictly on other consumers (*Carniv_obs_* = 0.038). This is in stark contrast to the null expectation that <2% of species are basal (*B_null_ =* 0.018), and the remaining are intermediate (*I_null_* = 0.977) with <1% TL2 herbivores (*TL2_null_* = 0.006). Moreover, the null model predicts one-third of species as carnivores (*Carniv_null_ =* 0.315), and the remaining two-thirds as omnivores (*Omni_null_ =* 0.662). However, our observation of almost no top consumers was not significantly different from the null expectation of the niche model (*T_obs_* = 3.85 x 10^-4^, *T_null_* = 0.006), and the fraction of cannibals was also similar to the null expectation (*Cannib_obs_* = 0.0312, *Cannib_null_* = 0.058).

Our web contains a higher fraction of herbivorous links (*HerbLink_obs_* = 0.135, *HerbLink_null_* = 0.018) – especially a higher fraction by TL2 herbivores – than expected (*TL2Link_obs_* = 0.071, *TL2Link_null_* = 4.87 x 10^-5^, Fig. 3F) and tends to be shorter in terms of mean and max trophic level (*meanTL_obs_* = 2.22, *meanTL_null_* = 4.37; *maxTL_obs_* = 4.62, *maxTL_null_* = 6.17). Additionally, the mean trophic level of top species in our web is 2 (i.e., herbivores), significantly lower than expected (*meanTLTop_obs_* = 2.0, *meanTLTop_null_* = 4.81). This may be a limitation of the dataset rather than a true signal; our temperate forest system does not have megaherbivores, and we expect most insect herbivores to experience parasitoidy or natural enemies in aboveground terrestrial habitats.

Our web exhibits a higher average generality among consumers (*meanGen_obs_* = 186.7, *meanGen_null_* = 143.5, Fig. 3H) and a greater variability in generality overall (*GenSD_obs_* = 2.82, *GenSD_null_* = 1.19, Fig. 3J). Indeed, the distributions of generality between our web and the simulated niche models are significantly different, both visually (Fig. 2B) and statistically according to Kolmogorov-Smirnov tests (see Supplementary Table 2). These deviations can be attributed to two properties: first, a small core of hyper-generalist omnivores (*meanGenOmniv_obs_* = 481.3, *meanGenOmniv_null_* = 144.4; *GenSDOmniv_obs_* = 1.35, *GenSDOmniv_null_* = 1.17); second, a long tail of TL2 herbivores with greater generality and variability of generality than expected by the niche model (*meanGenTL2_obs_* = 20.8, *meanGenTL2_null_* =1.42, Fig. 3I; *GenSDTL2_obs_* = 1.30, *GenSDTL2_null_* = 0.40, Fig. 3K). However, the TL2 herbivores produced by the niche model are far more specialized than the average consumers in our web (compare to *meanGen_obs_*).

**Figure 2.**
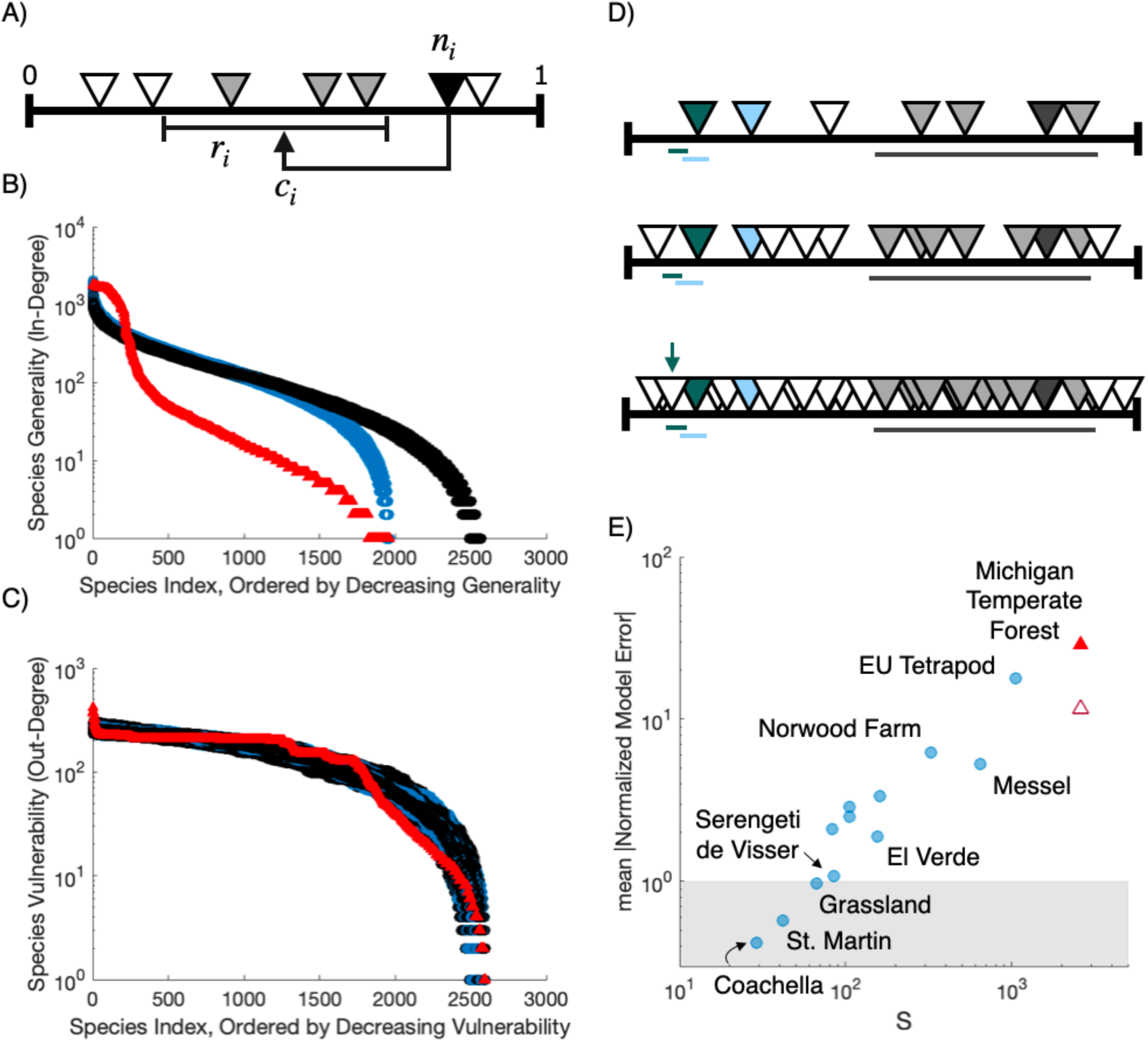
The Michigan Temperate Forest food web structure deviates from null expectations of the niche model due to problems of niche packing at high richness. **(A)** We use Williams & Martinez’ (2000) niche model to simulate an ensemble of null-model food webs with trophic species richness (*S*) and directed connectance (*C*) matched to the empirical food web. For each species *i*, the model randomly draws a niche value from a uniform distribution (*n_i_*∼*U*(0,1)) and assigns *i* a feeding range along the niche axis (*r_i_* = *n_i_x*, where *x* is β-distributed with *E[x]* = 2*C*) centered at a value (*c_i_*) below its niche value (*c_i_* ∈ [*r_i_*/2, *min*(*n_i_*, 1 − *r_i_*/2)]). In this way, the higher the niche value of a species, the wider its feeding range. Then, *i* feeds on all species *j* with niche values in that range or is considered a basal species if the range is empty. Only trophic species with a unique set of consumers and resources are permitted. To guarantee a basal species, the feeding range of the species with the lowest niche value is set to zero (*r_i_* ∶= 0). **(B)** The observed in-degree distributions for the Michigan Temperate Forest web (red triangles) compared to in-degree distributions from *N* = 1,000 simulated niche model webs with matched *S*,*C* (black circles) or matched *S*,*C*, and fraction of basal species, *B*, achieved by setting more species’ feeding ranges to zero, blue circles). Degree is on a log scale (*y*-axis), with species (*x*-axis) ordered by decreasing degree. In-degree is generality, the richness of diet. The empirical web has a core of high-degree generalists (birds and bats) and a long tail of lower-degree trophic-level 2 herbivores (insects), leading to a significantly different distribution of generality than predicted by the niche model. **(C)** Observed and simulated out-degree distributions. Out-degree is vulnerability, the richness of predators. **(D)** The strict one-dimensional hierarchy of the niche model does not accommodate the rich community of trophic-level 2 herbivores (*TL*2) observed in the empirical food web. As *S* increases, the niche axis becomes increasingly and uniformly packed with species, represented here as triangles. Horizontal bars represent the feeding ranges of each colored species. At high *S*, the chance that species will have empty feeding ranges decreases, decreasing *B* (e.g., the green species). The fraction of TL2 herbivores also decreases with *S* because herbivores’ feeding ranges must be perfectly placed to include a rare basal species but exclude any consumers. For example, the blue species feeds only on the green basal species but becomes a carnivore when the green species is switched to a consumer at high *S*. The fraction of top species (i.e., without predators, *T*) also decreases with *S* because the broad feeding ranges and tendency to cannibalism of high niche-value species increasingly cover the niche axis with predators and excludes those species from being top (e.g., the gray species). **(E)** Mean absolute normalized model errors (NMEs) across all traditional niche model properties. Formatting follows Fig. 3 but absolute rather than signed errors are shown, and points are on a log-log scale.

**Figure 3.**
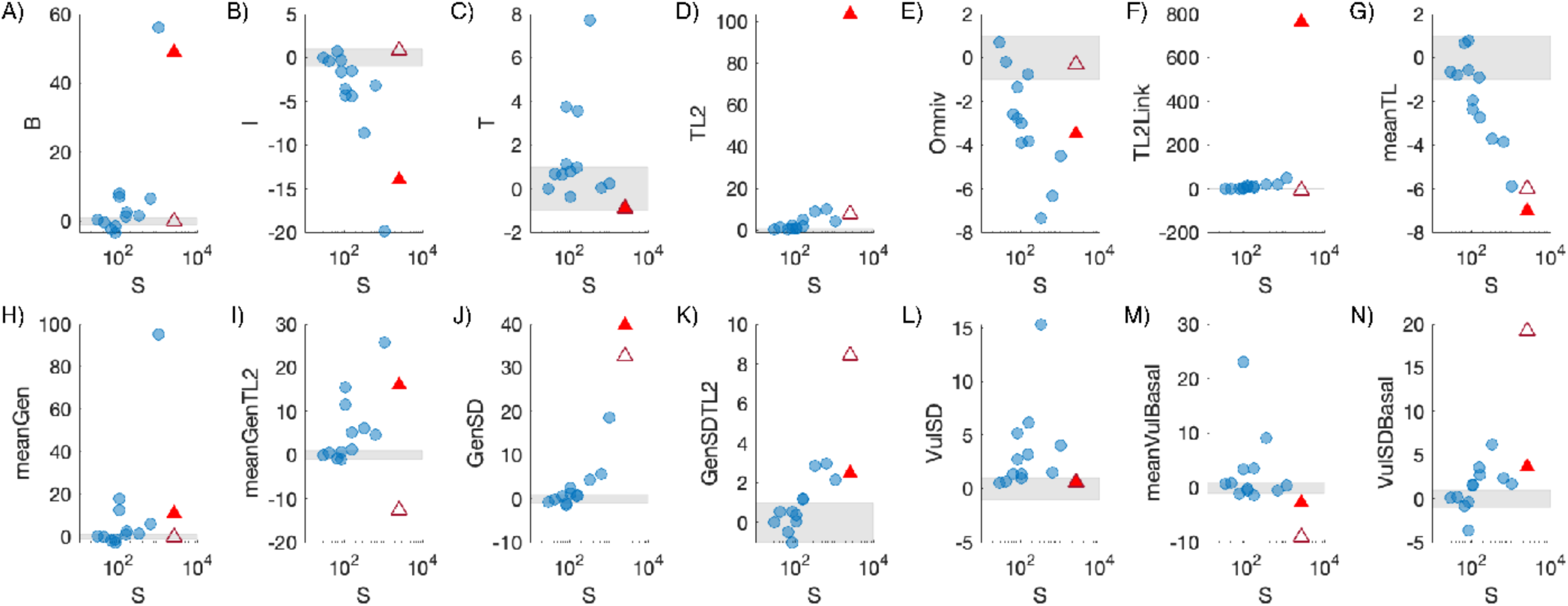
Structural properties of the Michigan Temperate Forest web deviate from previous aboveground terrestrial food webs and null model expectations. Points are normalized model errors (NMEs) calculated for empirical food webs compared to null expectations from an ensemble of *N* = 1,000 niche model food webs simulated with matched trophic species richness (*S*) and directed connectance (*C*). NMEs > 0 or < 0 indicate that observed properties are greater or less than expected, respectively. |NMEs| > 1 (outside of the gray box) are significantly different from null expectations at the 95% confidence level. Blue dots represent previously published food webs (Table 1). Red triangles are the Michigan temperate forest food web compared to the niche model with matched *S*, *C* (filled) or matched *S*, *C*, and *B*, the fraction of species with no resources (i.e., basal species, achieved by setting species’ feeding ranges to zero, hollow). Plots show that NMEs broadly scale with *S*, but the magnitudes of deviation associated with our web are greater than previously observed. See Methods for full definitions of food web properties. Fraction of **(A)** basal, **(B)** intermediate, **(C)** top, **(D)** strict herbivore (i.e., trophic level = 2), and **(E)** omnivorous species. **(F)** Fraction of feeding links by TL2 herbivores. **(G)** Average trophic level. Average generality of **(H)** consumers and **(I)** TL2 herbivores and normalized standard deviation of generality of **(J)** consumer species and **(K)** TL2 herbivores. Normalized standard deviation of vulnerability of **(L)** resource species and **(N)** basal species. **(M)** Average vulnerability of basal species.

Our web does not exhibit significant deviations in average vulnerability (*meanVul_obs_* = 141.1, *meanVul_null_* = 141.1) or variability of vulnerability (*VulSD_obs_* = 0.583, *VulSD_null_* = 0.583, Fig. 3L) compared to the null expectation, despite significantly different degree distributions (Fig. 2C). However, our web specifically has a lower vulnerability of basal trophic species than expected (*meanVulBasal_obs_* = 77.6, *meanVulBasal_null_* = 139.7, Fig. 3M) and a higher variability of such vulnerability (*VulSDBasal_obs_* = 1.052, *VulSDBasal_null_* = 0.580, Fig. 3N).

To investigate the extent to which deviations in the structure of our food web from both previous webs and null expectations can be attributed simply to differences in species composition, we also compared our web to an ensemble of niche models with a matched fraction of basal trophic species, *B* (Fig. 3A, hollow triangle). This correction automatically fixes *I* (Fig. 3B) and *meanGen* (Fig. 3H) by restricting the available links to the correct number of consumers (Fig. 2B), thereby improving *Omniv, TL2*, and *TL2Links* (Fig. 3D-G, J). However, this causes a compensatory overestimation of total herbivorous links, TL2 generality, and basal vulnerability (*HerbLink_nullB_* = 0.344; *meanGenTL2_nullB_* = 105.1, Fig. 3I; *meanVulBasal_nullB_* = 200.0, Fig. 3M), as well as a further underestimation of the variabilities of TL2 generality and basal vulnerability (*GenSDTL2_nullB_* = 0.92, Fig. 3K; *VulSDBasal_nullB_* = 0.31, Fig. 3N). We also tested whether the higher *B* and herbivory interactions in our original trophic-species web were attributable to the presence of basal animals. After removing these species and links, we recovered similar properties to the original web (*B_obs_* = 0.232, *I_obs_* = 0.768, *TL2_obs_* = 0.491, *HerbLink_obs_* = 0.117).

This finding clarifies why our web deviates from niche model predictions (Fig. 2D). The strict hierarchy of the niche model causes decreasing *B* and *T* with increasing *S* and therefore becomes very unlikely to generate TL2 herbivores. However, even if forced to generate larger *B*, TL2 herbivores remain rare, with low variability in generality. This is because TL2 herbivores must have feeding ranges that only include basal species, which is most likely with narrow ranges (and therefore low niche values). In contrast, the TL2 herbivores in our web are numerous, have diets on average wider and more variable in size than those produced by the niche model, and their trophic species niches are distinguished by highly specific herbivory interactions rather than by their predator interactions.

## 4) Discussion

The food web for the Michigan Temperate Forest (MTF) presented here is the largest yet published (Table 1, [6,10]) and begins to shine light on the remarkable richness of feeding interactions between plants and animals in aboveground terrestrial systems. Our cumulative approach allowed us to evenly resolve the diets of both vertebrate and insect feeding guilds, revealing a clear pattern among herbivory and carnivory interactions. Herbivory interactions are far rarer (<15% of trophic links) than carnivory interactions, but the former were primarily sourced from the largest guild in the web (∼50% of trophic species) – the strict (i.e., trophic-level two) insect herbivores with variably-sized but highly specific diets, both taxonomically and in terms of plant tissue types. In addition, carnivory interactions dominate the web (>85% of trophic links) but were primarily sourced from a small fraction of trophic species (<4.6%) – hyper-generalist birds and bats thought to feed opportunistically on entire insect orders. The combination of these two properties in our food web leads to a structure that qualitatively deviates from previous aboveground terrestrial food webs (ATFWs) of lower taxonomic and trophic resolution, as well as the scale-dependent null expectations of the niche model.

Given the richness and complexity of our trophic web, the niche model predicted an even larger fraction of carnivory interactions (>98%), stemming from the minimal fraction of basal trophic species with no consumers (*B* < 2%) and herbivores (*TL2* < 1%), ultimately resulting in an overall normalized model error far greater than for any previous webs studied here (MTF mean |*NME*| = 28.9, Fig. 2E). ATFWs using the classic ‘lumping’ approach (Coachella Valley, St. Martin Island, El Verde Rainforest, Serengeti de Visser [50]) are generally small, with low *B* and *TL2* (due to extremely coarse representation of plants and insects) and high fractions of carnivory links (due to the disproportionate resolution of vertebrate predators). These properties align with the predictions of the niche model, as quantified by normalized model errors (0.42 < mean |NME| < 1.88, Fig. 2E). High-resolution source and sink ATFWs (Scotch Broom [51], UK Grassland [52,53], High Arctic [38,54–57], Norwood Farm [36]) record primarily parasitoids and parasites as the higher trophic-level consumers but generally include herbivory links on multiple plant tissues. The relative lack of generalist or opportunistic predators leads to high fractions of top species and very low omnivory, both in contrast to our web and the expectations of the niche model (0.97 < mean |NME| < 6.21). Most comparable to the MTF in terms of size and resolution is perhaps Messel Forest [41], though it is from the early Eocene and contains many now extinct species. While the MTF has greater *TL2*, omnivores, and carnivory links, as well as a lower fraction of herbivory links, the significantly different distributions of generality are likely the greatest contrast (Supplementary Table 2). Messel Forest has fewer TL2 herbivores with narrower and less variable diets (deviating less from niche model predictions) and does not show an inflection in its (log) degree distribution caused by a group of hyper-generalists. As a result, the Messel Forest web shows an overall better correspondence to null expectations of the niche model (mean |NME| = 5.28), despite a substantially larger richness than classic webs.

The niche model is considered to successfully reiterate the properties of natural food webs and is even used to simulate network structures for studies of food web dynamics, though it is known to underestimate *TL2* and overestimate average trophic level [21,44]. This success can broadly be attributed to two mathematical properties, shared by other generating models of food web structure [1,3,58–60]: first, that species can be strictly ordered along a one-dimensional axis by their niche values (*n*_1_ < *n*_2_ < ⋯ *n*_2_), and second, that species’ feeding ranges (*r_i_*) are exponentially-distributed, with a decaying probability of feeding on species of lower niche values. The superior performance of the niche model (given its simplicity) is explained by its third property [8]: it generates “interval” webs, because species feed on all resources within intervals of the niche axis (i.e., they have contiguous diets). These properties are also ecological hypotheses for mechanisms that structure food webs [40]. For the MTF, the largest magnitudes of deviations (|NME| > 95) were in terms of species composition, which we attribute to issues with niche packing (Fig. 2D, but see [61]). This suggests that the MTF does not satisfy the first or third properties, since allowing more herbivores on a packed niche axis would require allowing an additional axis of food selection among herbivores of the same niche value or breaks in their diet contiguity. In fact, the MTF, like many others, is not strictly interval. As such, the more pertinent question is the level of intervality of the MTF relative to other webs of its scale and whether this could cause the large magnitude of deviations we observed. Unfortunately, this is combinatorically intractable for us to assess using current methods [3,62]. Lastly, the MTF does not satisfy the second property of exponentially distributed feeding ranges (Fig. 2B-C). We hypothesize that for the niche model to better reproduce MTF properties, links would need to be preferentially allocated to a small group of generalist species and another axis associated with eating plants would need to be introduced to allow for rich communities of herbivores with variably-sized diets [7].

Whether the structural patterns for the MTF could be general among aboveground terrestrial ecosystems remains to be tested. We observed an overall increase in NMEs with richness (Fig. 2E, Fig. 3), such that we cannot confidently reject the null hypothesis that our observed structure can be attributed to scale-dependent errors associated with the niche model. However, the MTF appears to exhibit a qualitatively different structure than previous works, including a potentially novel degree distribution. Kolmogorov-Smirnov tests rejected the hypothesis that degree distributions of generality and vulnerability for our web were sampled from the same underlying process as the distributions of each of the other ATFWs studied here (p < 1.0 x 10^-5^ in all cases, Supplementary Table 2). This could imply that terrestrial food webs do have unique structure – potentially driven by the opportunism of predators and a lack of hierarchy among strict herbivores – which may only be observed in the context of high taxonomic and trophic resolution among a rich community of plants, insects, and vertebrates, such as in our temperate forest system.

As always, there are caveats to these conclusions, primarily associated with our methodological approach and general data limitations. It is possible that the critical hyper-generality of bats and birds in the MTF can be attributed simply to lack of taxonomic resolution regarding their specific foraging preferences (e.g., microlepidopterans, a paraphyletic group with <20 mm wingspans, may be too small to be eaten by vertebrate predators). Yet studies examining species-specific diets of bats and birds have supported hyper-generalism (e.g., [63]), even to the extent that molecular identification of species in bat guano presents a roughly equivalent snapshot of insect biodiversity as traditional blacklight sampling of insects [64]. We also know that many plant and insect species are missing interactions because we were not able to find or verify species-specific data (due to taxonomic or other data limitations) or because records were too vague (e.g., “eating seeds” without further specificity). It is therefore possible that herbivory links were underrepresented relative to carnivory in our observations. Interactions could be more thoroughly refined to account for species’ traits and ecological habits, which would likely convert some links currently considered plausible to effectively ‘forbidden links’ *sensu* Jordano et al. [65,66]. Nevertheless, opportunistic carnivory links would still dominate the web and additional herbivory links would likely be taxonomically specific and trophically distinct, which may even serve to increase trophic species richness by distinguishing plants’ or herbivores’ trophic niches. In short, a further refinement of feeding interactions would most likely align with the food web structure described here.

Compiling taxonomic and feeding records for diverse groups into cumulative food webs has the caveat of potentially introducing diverse and compounding sampling biases, making it difficult to quantify uncertainty in the reported food web structure [16]. In the MTF, for example, the relative representation of some groups can be assessed directly through rarefaction curves over field seasons (e.g., lepidopteran leafminers), while some groups (e.g., non-insect arthropods) include diversity for which we were not able to source even regional lists as reference points (e.g., acariform mites). Likewise, feeding on some types of resources, especially by specific taxonomic groups, is more readily observed or charismatic than others, and therefore better documented. As food web research moves towards synthesizing bigger and more diverse data, an important theoretical question for future work is to develop quantitative methods for characterizing how uncertainty may affect observed network structure (see [16,67] for promising methods).

Like previous researchers, we chose to limit our scope to the aboveground portion of our food web. However, we recognize that all ATFWs are intimately and inextricably coupled with belowground soil food webs. A substantial fraction of plant biomass may exist belowground [68], creating habitat structure, and providing food for organisms that consume roots, exudates, and detritus. Decomposers return nutrients generated from aboveground waste to the soil for reuptake by plants, often facilitated by mutualistic symbionts (e.g., mycorrhizae) [69]. Moreover, many consumer species (including some in the MTF) live or feed belowground during certain life stages or times of year. As such, complex food web dynamics belowground have considerable impacts on food web dynamics aboveground, and interactions between these two habitats can significantly influence ecosystem-level processes [70,71]. While this fact has long been recognized as a bias in ATFW research, rarely are above-and belowground food webs recorded in the same system (likely due to the logistical challenges of sampling belowground). Doing so represents a critical research frontier for terrestrial food webs as we seek to understand their structure, dynamics, and emergent ecosystem functions.

This study is only our first step towards documenting the immense taxonomic diversity and trophic complexity in the temperate forests of University of Michigan Biological Station (UMBS). Ecological networks have historically been published and analyzed as static structures, encapsulating the biases and practical limitations of their collection. As such, publication in online databases and consistent re-use in meta-analyses by ecologists and network scientists can perpetuate errors [17,72]. Our ability to create a large, highly resolved, expert-vetted ATFW was made possible through decades of research and observations at the University of Michigan Biological Station and highlights the importance of local knowledge, taxon-specific expertise, and collaboration among scientists, students, and members of the public. As insights about the organisms present at UMBS will undoubtedly continue to grow, we consider the Michigan Temperate Forest web to be a living dataset that can be revised and expanded through time. To that end, our database is publicly available [43], and we are soliciting revisions, corrections, and additions that will allow its continual improvement.

## Acknowledgements

We thank Adam Schubel, Jason Tallant, Aimee Classen, Knute Nadelhoffer, and other current and former University of Michigan BioStation (UMBS) staff for providing the original species lists and hosting the living version of the dataset. We are immensely grateful to Teresa Pegan, Eric Gulson, Simone Oliphant, Nate Sanders, Daniel Swanson, Anton Reznicek, Erika Tucker, and undergraduate research team Kathrine Northman, Taylor Brubaker, John Kelly, Lynnae Gilman, Matthew Palumbo, Lex Newman, and Stephan Verral for contributions to data acquisition and vetting. We acknowledge that the Indian Point Reserve (gifted to UMBS in 1987) includes lands of the Burt Lake Band of Ottawa and Chippewa Native Americans from which they were brutally and illegally evicted in 1900. The University of Michigan Undergraduate Research Opportunities Program (UROP) paid the undergraduate researchers for their time. This work was partially funded by NSF grants DEB-2129757 and DEB-2224915 to F.S.V.

## Data Availability Statement

Species and interaction data are openly available via the Environmental Data Initiative at https://doi.org/10.6073/pasta/840d70788bde4692a7d6d45f8d04376f. Electronic Supplementary Information is available online.

## Electronic Supplementary Information

### Supplementary Methods

#### i) Site description

The University of Michigan Biological Station (UMBS) includes ∼10,000 acres of land that is used for teaching and research in northern lower Michigan, USA (45°35.5′ N, 84°43′ W). The property is composed predominantly of dry-mesic, northern hardwood forests with patches of wooded wetlands (hardwood conifer swamp) flanked by two lakes [42]. This is a strongly seasonal, temperate system with historically cold, snowy winters (average minimum temperature of –12.1°C in January) and hot, humid summers (average maximum temperature of 26.2°C in July; [77]). Soil composition varies across microhabitat patches from sandy outwash plain to moraine. Prior to acquisition by the University of Michigan in 1909, the landscape was almost completely logged and burned, with only a few old-growth forest tracts remaining. Therefore, the vegetative landscape is relatively uniform in age, with variation in stand structure and composition attributable to glacial landforms and differences in soil composition [78,79].

UMBS is contiguous to other forested habitats, allowing free movement in and out for mobile species, including those that seasonally migrate. Such transient organisms, coupled with those that are only active or present aboveground during certain times of year, lead to strong effects of seasonality on species composition. Though their impacts may only be temporary, we included these species in our analyses because they may represent critical consumers or resources for local species at certain times of year. In particular, most species of ectotherms at UMBS are inactive during winter months, and many migratory bird species are present only during the spring and summer. By pooling observations into a cumulative web, we avoid sensitivity of food web structure to this spatial and temporal transience.

#### i) Species list

To construct the food web, we began from taxonomic lists of mammals, amphibians, reptiles, vascular plants, birds, insects, and non-insect arthropods. These lists represent an accumulation of decades of records at UMBS, from resident biologists’ personal observations, student papers and research projects, regional lists, museum specimens, online databases (such as eBird and iNaturalist), and semi-regular BioBlitz events, in which teams of biologists roamed the site and identified as many organisms as possible.

Where possible, we updated records to the most recently accepted species names according to the Integrated Taxonomic Information System (ITIS) using the “taxize” package in R [80]. We excluded taxa that could not be resolved to at least genus-level (e.g., parasitoid wasp family Diapriidae). For taxa that could only be resolved to genus, we excluded those that are highly speciose in the Nearctic (> 20 species, e.g., the bloodworm genus *Chironomus*) and those that had congenerics already included in the food web (e.g., six species of masked bee *Hylaeus* were included, but another unidentified species was excluded). Finally, we treated variants and subspecies (e.g., deer mouse *Peromyscus maniculatus gracilis*) as their binomial species name. Hereafter, we refer to all taxa occurring at UMBS as “species,” though a small fraction (4.5%) are genera.

The species lists were vetted and approved by experts (generally, the authors) that have extensive knowledge of the communities and natural histories of the organisms that occur in the region. In determining which species to include or exclude, we excluded any species that do not occur at UMBS or do not have a significant lifestage or feeding behavior in aboveground terrestrial habitats [76]. We defined “aboveground terrestrial habitats” as land above the soil layer (including leaf litter and above), on terrestrial plants (growing potentially over but not exclusively in water), or in air (above other terrestrial habitats). Thus, we excluded species that live and feed primarily on the surface of aquatic habitats, such as waterlilies (Nymphaceae) and water bugs (Belostomatidae). Additionally, given our primary focus on even resolution among aboveground plants, insects, and vertebrates, we chose to exclude some major groups, such as bryophytes, fungi, lichens, and molluscs.

#### ii) Feeding interactions

To assemble a list of potential feeding interactions between species, we referenced region-specific field guides, online databases (e.g., Animal Diversity Web, Birds of the World), and central repositories that automatically scrape data from museum records and the web (e.g., Global Biotic Interactions). Trained undergraduate researchers searched sources by species name and recorded predators, diet, and other potential feeding interactions with as much resolution as possible. Each focal taxon was resolved to species-level, but their interaction partners could be recorded at any taxonomic level (e.g., species *x* eats Family *y*). If we could not find information on a given focal species, we did not extrapolate from closely related species. We included all records of direct interactions among species in our system with a bioenergetic flow (i.e., one species consuming another), regardless of lifestage or potential ecological effects (i.e., whether potentially “mutualistic” or “antagonistic”, see the recent framework for studying aboveground terrestrial food webs, [14]).

Our records were supplemented, revised, and annotated by the same experts who vetted the species lists. Where possible, records were assigned to experts for both the focal species and its partner’s group (that is, two separate experts). We provide the final list of annotated feeding records and their experts online [43].

Experts broadly categorized records by their focal resource type as animal tissues, both (1) live tissues and as prey, and (2) scavenged as carrion, carcasses, or other decaying animal remains, and plant tissues, grouped as (3) leaves and stems, (4) flowers, nectar, pollen, etc., (5) seeds, fruits, etc., and (6) wood and bark. Hereafter, we refer to these resource types simply as “prey,” “carrion,” “leaves,” “flowers,” “seeds,” and “wood.” We excluded records of detritus resources that could not be resolved taxonomically.

Experts were asked to assess the plausibility of the recorded interactions occurring at UMBS. Interactions between species *x* and *y* were considered plausible if the two species co-occur (with respect to phenology, activity patterns, microhabitat usage) and have no trait incompatibilities (it would not be possible for *x* to acquire, ingest, or assimilate *y*). Interactions were considered plausible even if they could be considered inefficient (*x* can consume *y*, but this is rare or unlikely, especially if more rewarding and easily acquired foods than *y* are available). Interactions that are non-consumptive (e.g., nesting or hunting locations) or occur outside of aboveground terrestrial habitats were excluded even if they occur between local species (e.g., least sandpiper *Calidris minutilla* preys upon toad *Anaxyrus americanus* tadpoles, but this occurs exclusively in aquatic habitats).

Experts were also asked to assess the plausibility of the taxonomic level at which interactions were recorded. For example, when Wilson’s warbler (*Cardellina pusilla*) is recorded to eat beetles (order Coleoptera), it is unclear if: (case 1) *C. pusilla* eats every species of Coleoptera, potentially or opportunistically, including all local Coleopterans, or (case 2) *C. pusilla* eats (at least) one species of Coleoptera that was not identified in the original record and which may or may not be local. Either case may be possible, depending on the biology of the species. At one extreme (case 1), the lack of taxonomic resolution in the record could reflect an ecologically relevant lack of discrimination by the focal species (e.g., opportunism due to sensory capabilities in distinguishing between predators or resources), with the probability of interactions occurring between any given partner species determined broadly by their abundances. At the other extreme (case 2), the lack of taxonomic resolution could simply reflect a lack of knowledge, representing a summary of potentially highly specific interactions. In the first case, experts accepted the interaction, while in the second, experts rejected the interaction unless it could be plausibly revised for local taxa.

In the original sources, interactions were often recorded following traditional disciplinary categorizations. In the absence of other information, we assumed that animals recorded as “eating,” “feeding on,” “consuming,” “parasitizing,” or “hosted by” plants were feeding on leaves. We assumed that feeding on wood or bark was specifically recorded as such. We assumed that animals recorded as “pollinator of,” “visiting flowers of,” “visiting,” or “mutualist of” plants were feeding on flowers. Other mutualisms involving potential feeding such as seed dispersal (through frugivory, scatterhoarding, etc.) and ant protection (of hemipterans or plants) were noticeably underrepresented in our records. We assumed animals recorded as “disperser of” plants were feeding on seeds. We assumed that interactions between animals recorded as “kills,” “predates upon,” “host of,” “parasite of,” or “parasitoid of” involved consumption of animal tissue, and that “scavenges” reflected feeding on carrion. When necessary, these default assumptions were revised by experts to reflect true feeding on another resource type or a lack of feeding altogether.

#### iii) Sampling biases

Vertebrates and vascular plants are likely well-sampled in our system. Butterflies and moths are reasonably well-represented as they have been the focus of repeated target sampling. For some groups of insects (especially beetles, wasps, and true bugs), regional datasets suggest that as many as a few thousand more species occur but currently remain undetected at UMBS. Broadly, the species most likely missing from our list are those that are specialists, cryptic, and/or rare. Regional datasets also suggest that we are missing a smaller number of predatory non-insect arthropods (including up to 200 spp. of spiders, harvestmen, centipedes, millipedes, and terrestrial isopods). Among the major insect groups, true flies are likely the most underrepresented. This is not necessarily surprising as flies are extremely ecologically diverse and require specialized collecting – groups such as the fungus-associated flies (a notable exclusion in our list) would not necessarily be detected in malaise traps. We are also aware we are missing potentially many species of groups such as acariform mites, which were both underrepresented in our system and for which we did not have a regional reference list. Finally, for the purposes of limiting the scope of our work, we excluded microbes, fungi, detritus, non-vascular plants (including up to 900 spp. of mosses, liverworts, and lichens), and many non-insect invertebrate groups (including up to 100 spp. of gastropods) from our original species list and focused only on the species interactions that occur primarily in aboveground terrestrial habitats.

#### iv) Food web structure

To simulate niche model food webs with matching *S* and *C* to the empirical food webs with low connectance, we seeded the algorithm with *S_seed_ = S +* 4 for the High Arctic, UK Grassland, Scotch Broom, Shortgrass Prairie, and Messel Forest webs, and *S_seed_ = S* + 10, *C_seed_* = *C* x 1.01 for the Norwood Farm web. At each replication, we aggregated the simulated web to trophic species and discarded it if it did not match the target *S*, *C*. For the purposes of discussion, we followed previous works to summarize deviations from null expectations as a composite *mean absolute Normalized Model Error* (|NME|) of traditional food web properties. We calculated *mean* |NME| as the average of the absolute value for the NME of each of the following properties: *Top (T)*, *Intermediate (I), Basal (B), Cannib, Herbiv (TL2), TrophicOmniv, LinkSD, GenSD, VulSD, PathLen, meanTL, MaxSim.* See full definitions in Supplementary Table 1, following [8,26,39,44–47]. Notice the nontraditional notation here and in the main text to accommodate new and additional focal properties.

### Supplementary Data & Code

taxonomic_web_analysis_EDI.m is a MATLAB script that builds and analyzes the taxonomic version of the Michigan Temperate Forest (MTF) network, using multiplex network and food web representations for different focal levels of taxonomic resolution.

simulated_matching_niche_EDI.m is a MATLAB script that runs the leave-one-out analysis on the MTF network, creates trophic-species food web representations of historical empirical aboveground terrestrial food webs (including the MTF), simulates matching niche models, analyzes their properties, and reproduces the main text figures.

**Supplementary Table 1.**
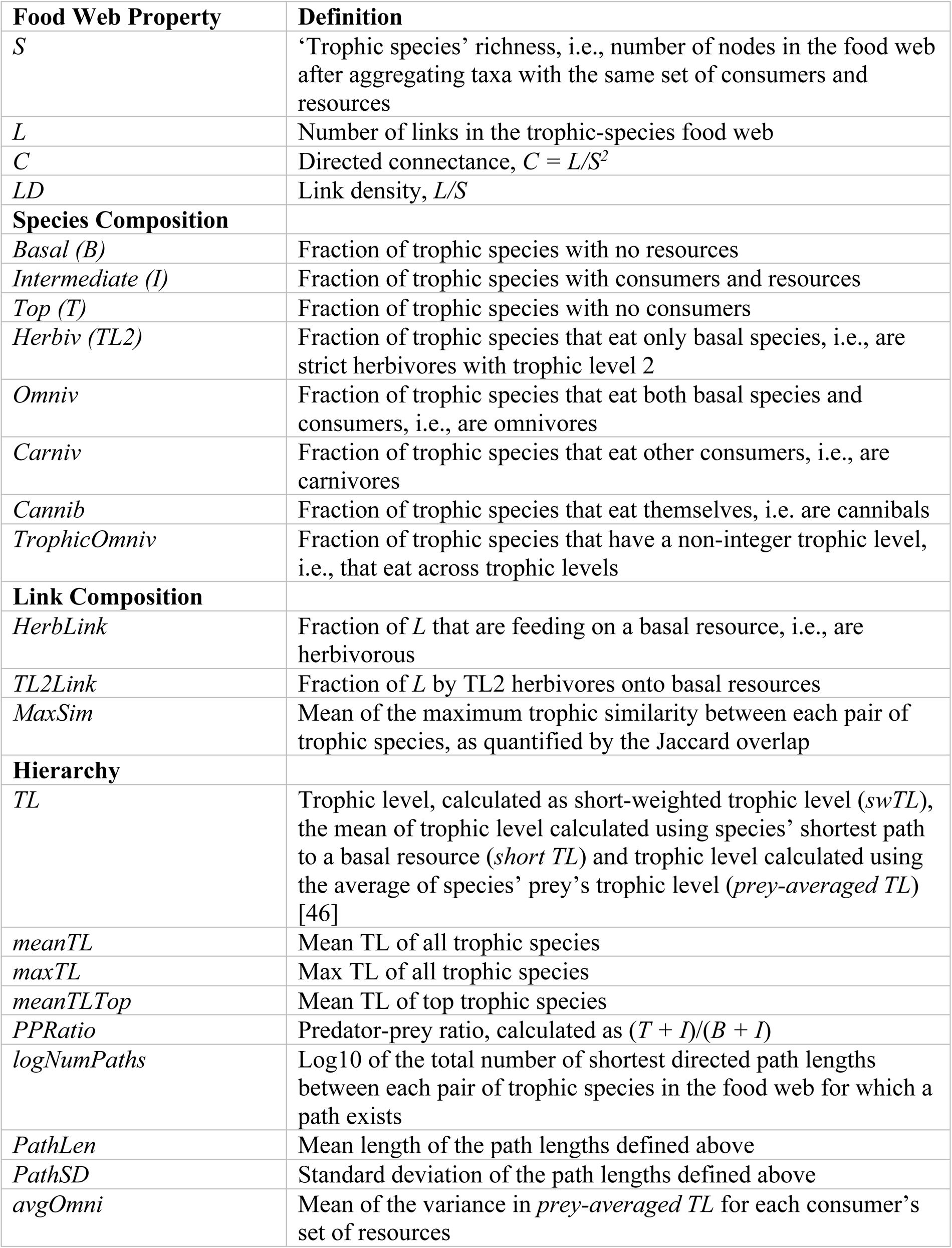
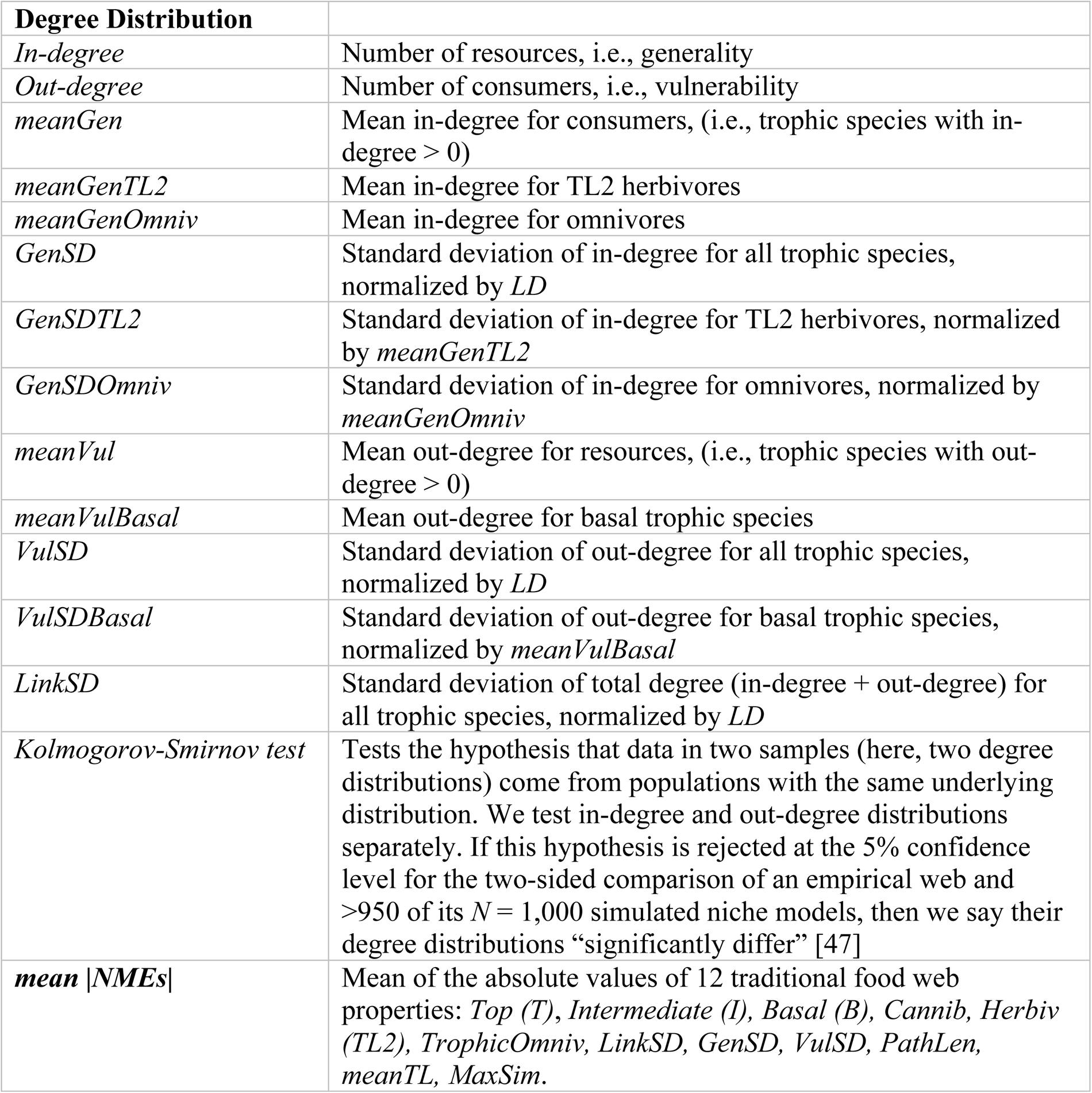
Full definitions of food web properties.

**Supplementary Table 2 Normalized model errors and Kolmogorov-Smirnov test results for empirical food webs**

**Supplementary Figure 1.**
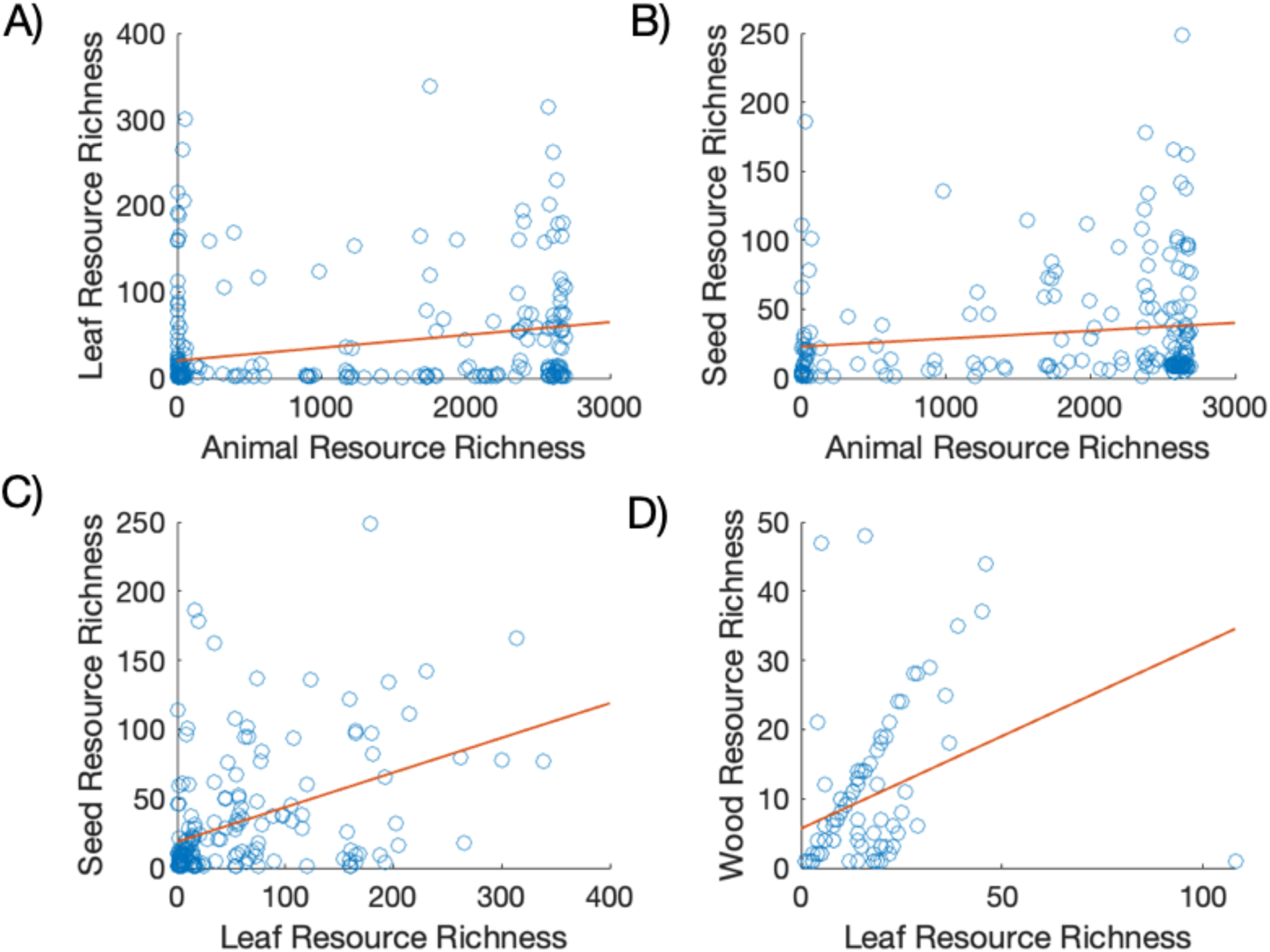
Correlations between consumer diet breadths feeding on different resources. Scatter plots showing the relationship between consumers’ diet breadths in terms of number of resource species when feeding on **(A)** animals and plant leaves (r = 0.26, p = 2.7 x 10^-6^, N = 319), **(B)** animals and plant seeds (r = 0.14, p = 0.044, N = 202), **(C)** plant leaves and seeds (r = 0.42, p = 7.2 x 10^-9^, N = 178), and **(D)** plant leaves and wood (r = 0.24, p = 0.017, N = 98). Each point is a species that feeds on both focal resources. Lines are least squares fits. Only relationships with significant Pearson’s correlations (r) are shown.

**Supplementary Figure 2.**
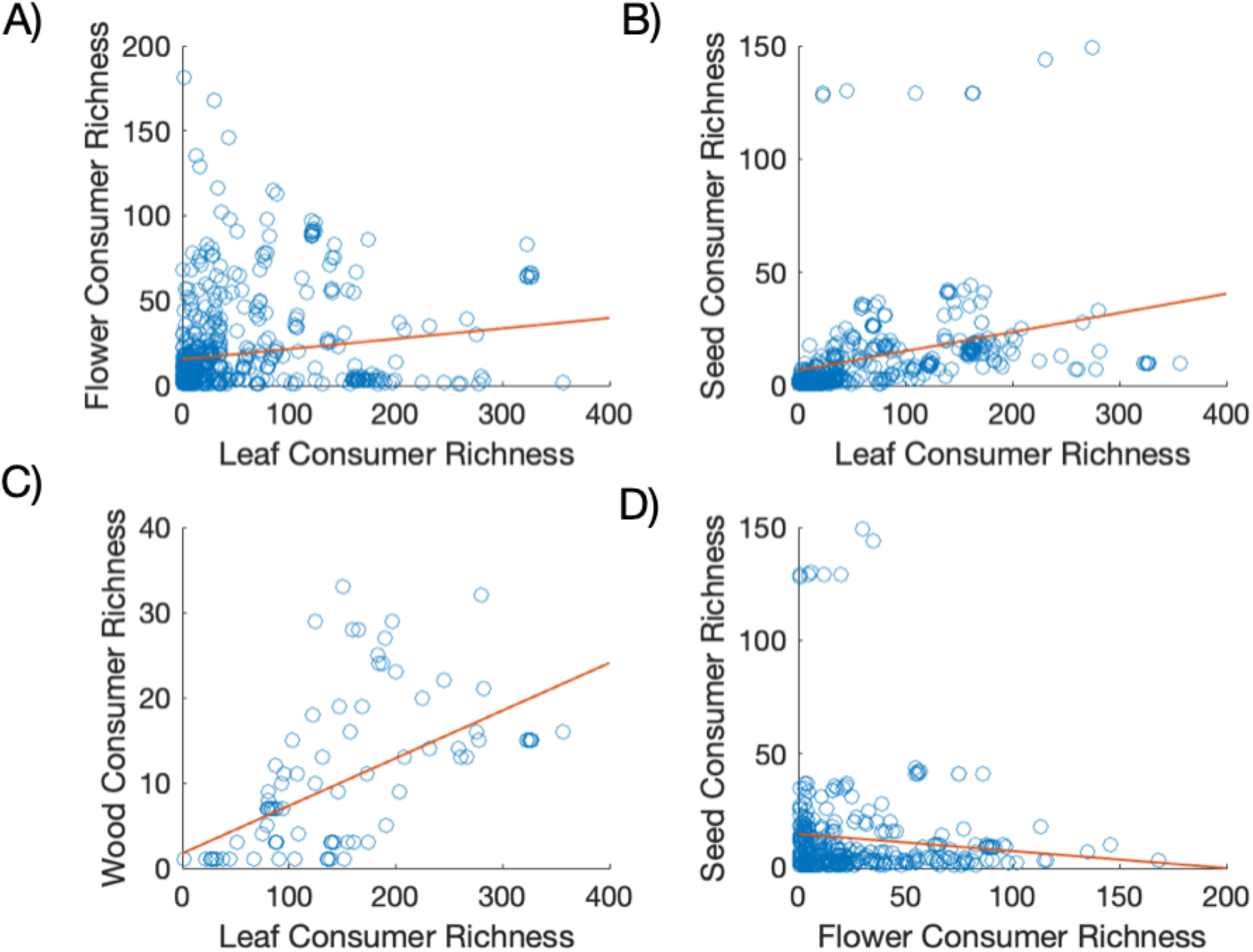
Correlations between animal richnesses hosted by plants on different tissues Scatter plots showing the relationship between the richness in terms of number of consumers species when feeding on plant species’. **(A)** leaves and flowers (r = 0.17, p = 1.8 x 10^-5^, N = 649), leaves and seeds (r = 0.35, p = 8.1 x 10^-17^, N = 548), **(C)** leaves and wood (r = 0.61, p = 1.7 x 10^-11^, N = 98), and **(D)** flowers and seeds (r = –0.11, p = 0.013, N = 510). Each point is a plant species with consumers that feed on both focal resources. Lines are least squares fits. Only relationships with significant Pearson’s correlations (r) are shown. This shows that the plants most consumed by leaf-herbivores are also most likely to host rich communities of consumers eating their other tissues. But there may be a trade-off in plants’ ability to support flower and seed eaters (potential pollinator and seed dispersal mutualists, respectively) or a deterrence effect between seed-and flower-eaters in our system (primarily birds and insects, respectively).

### Lay Summary

Terrestrial ecosystems are immensely complex, including diverse species feeding in diverse ways. Plants and insects engage in highly-specific interactions that have often been excluded from terrestrial food webs due to limited sampling and expertise. Here, we use records from a biological research station accumulated over ∼100 years to construct a food web for a Michigan temperate forest. We report ∼580,000 interactions among ∼3,800 species in a living dataset that is openly available to be supplemented and revised. This is the largest and most detailed food web yet reported and provides a valuable tool for ecosystem management and research.

