## Supplementary figures and images for "A highly resolved network reveals the role of terrestrial herbivory in structuring aboveground food webs"

### Supplementary Figure 1

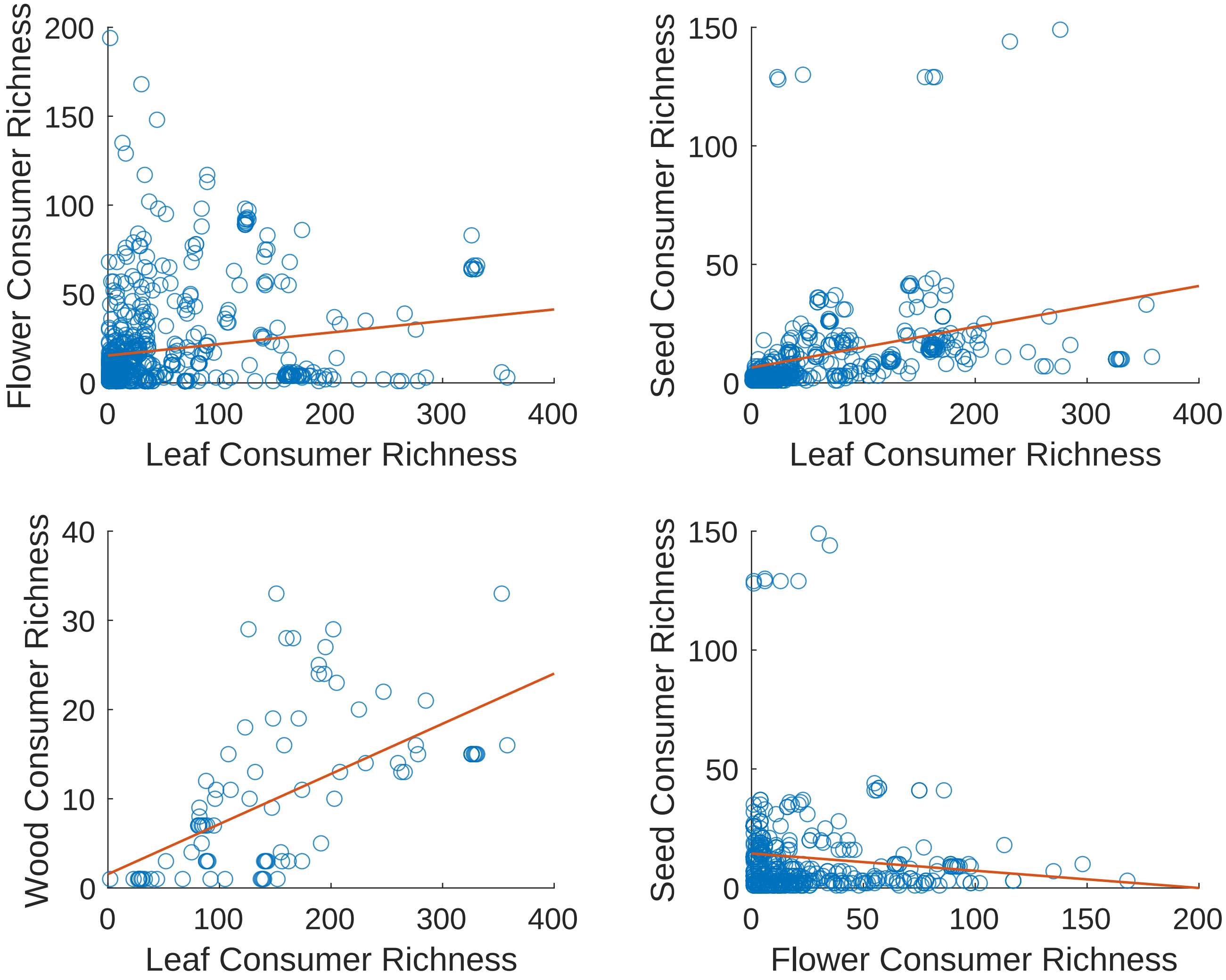

### Supplementary Figure 2

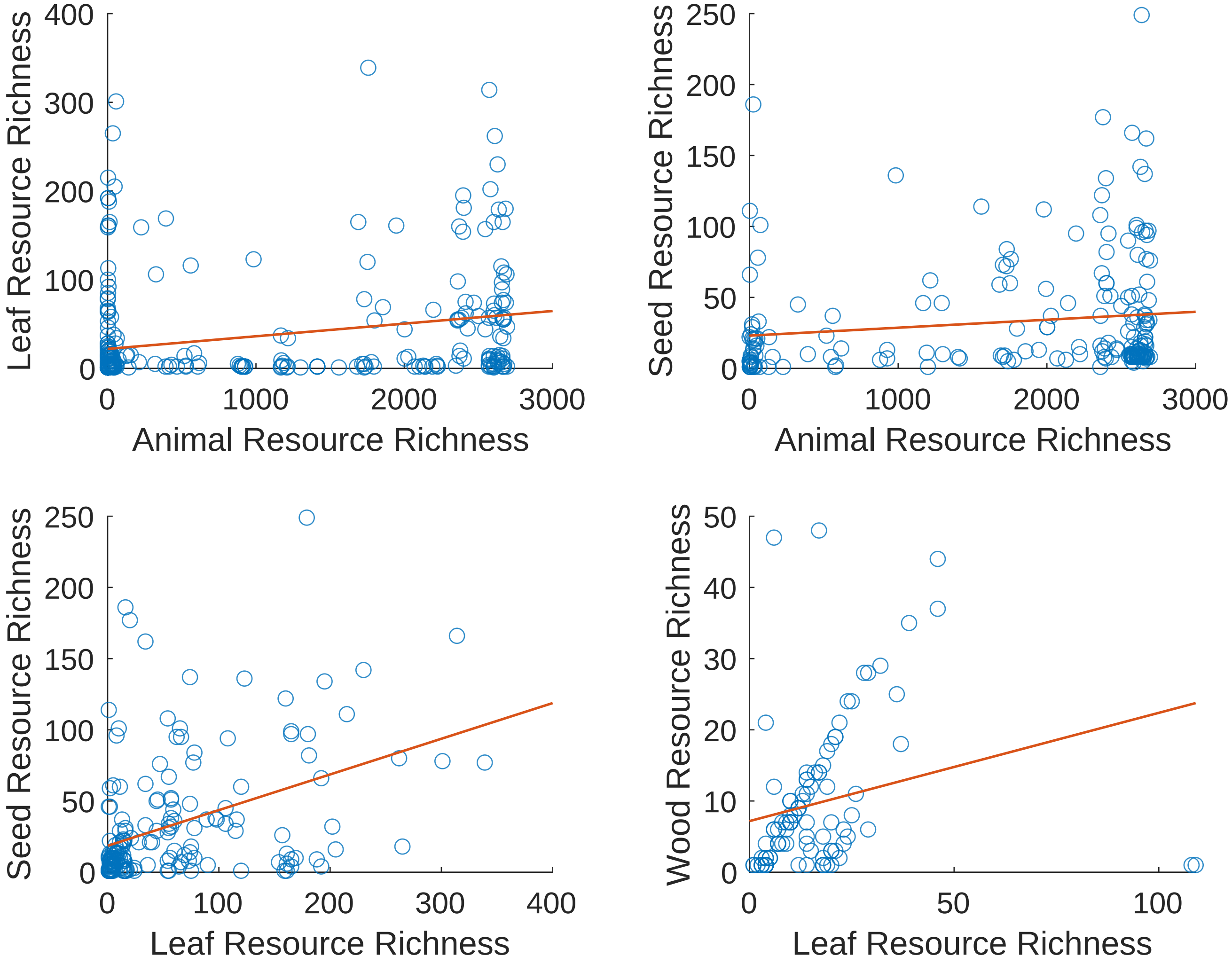
