## Supplementary Table 1 for "A highly resolved network reveals the role of terrestrial herbivory in structuring aboveground food webs"

**Supplementary Table 1 Full definitions of food web properties.**

| **Food Web Property** | **Definition** |
| --- | --- |
| *S* | ‘Trophic species’ richness, i.e., number of nodes in the food web after aggregating taxa with the same set of consumers and resources |
| *L* | Number of links in the trophic species food web |
| *C* | Directed connectance, *C = L/S^2^* |
| *LD* | Link density, *L/S* |
| **Species Composition** |  |
| *Basal (B)* | Fraction of trophic species with no resources |
| *Intermediate (I)* | Fraction of trophic species with consumers and resources |
| *Top (T)* | Fraction of trophic species with no consumers |
| *Herbiv (TL2)* | Fraction of trophic species that eat only basal species, i.e., are strict herbivores with trophic level 2 |
| *Omniv* | Fraction of trophic species that eat both basal species and consumers, i.e., are omnivores |
| *Carniv* | Fraction of trophic species that eat other consumers, i.e., are carnivores |
| *Cannib* | Fraction of trophic species that eat themselves, i.e. are cannibals |
| *TrophicOmniv* | Fraction of trophic species that have a non-integer trophic level, i.e., that eat across trophic levels |
| **Link Composition** |  |
| *HerbLink* | Fraction of *L* that are feeding on a basal resource, i.e., are herbivorous |
| *TL2Link* | Fraction of *L* by TL2 herbivores onto basal resources |
| *MaxSim* | Mean of the maximum trophic similarity between each pair of trophic species, as quantified by the Jaccard overlap |
| **Hierarchy** |  |
| *TL* | Trophic level, calculated as short-weighted trophic level (*swTL*), the mean of trophic level calculated using species’ shortest path to a basal resource (*short TL*) and trophic level calculated using the average of species’ prey’s trophic level (*prey-averaged TL*) [44] |
| *meanTL* | Mean TL of all trophic species |
| *maxTL* | Max TL of all trophic species |
| *meanTLTop* | Mean TL of top trophic species |
| *PPRatio* | Predator-prey ratio, calculated as (*T + I*)/(*B + I*) |
| *logNumPaths* | Log10 of the total number of shortest directed path lengths between each pair of trophic species in the food web for which a path exists |
| *PathLen* | Mean length of the path lengths defined above |
| *PathSD* | Standard deviation of the path lengths defined above |
| *avgOmni* | Mean of the variance in *prey-averaged TL* for each consumer’s set of resources |
| **Degree Distribution** |  |
| *In-degree* | Number of resources, i.e., generality |
| *Out-degree* | Number of consumers, i.e., vulnerability |
| *meanGen* | Mean in-degree for consumers, (i.e., trophic species with in-degree > 0) |
| *meanGenTL2* | Mean in-degree for TL2 herbivores |
| *meanGenOmniv* | Mean in-degree for omnivores |
| *GenSD* | Standard deviation of in-degree for all trophic species, normalized by *LD* |
| *GenSDTL2* | Standard deviation of in-degree for TL2 herbivores, normalized by *meanGenTL2* |
| *GenSDOmniv* | Standard deviation of in-degree for omnivores, normalized by *meanGenOmniv* |
| *meanVul* | Mean out-degree for resources, (i.e., trophic species with out-degree > 0) |
| *meanVulBasal* | Mean out-degree for basal trophic species |
| *VulSD* | Standard deviation of out-degree for all trophic species, normalized by *LD* |
| *VulSDBasal* | Standard deviation of out-degree for basal trophic species, normalized by *meanVulBasal* |
| *LinkSD* | Standard deviation of total degree (in-degree + out-degree) for all trophic species, normalized by *LD* |
| *Kolmogorov-Smirnov test* | Tests the hypothesis that data in two samples (here, two degree distributions) come from populations with the same underlying distribution. We test in-degree and out-degree distributions separately. If this hypothesis is rejected at the 5% confidence level for the two-sided comparison of an empirical web and >950 of its *N* = 1,000 simulated niche models, then we say their degree distributions “significantly differ” [45] |
| ***mean \|NMEs\|*** | Mean of the absolute values of 12 traditional food web properties: *Top (T)*, *Intermediate (I), Basal (B), Cannib, Herbiv (TL2), TrophicOmniv, LinkSD, GenSD, VulSD, PathLen, meanTL, MaxSim*. |
