## Supplementary material for "A highly resolved network reveals the role of terrestrial herbivory in structuring aboveground food webs": Data and Code: simulated_matching_niche_EDI.m

```
% %% LEAVE ONE OUT ANALYSIS
% load('multiplex_network.mat')
% 
% L = size(multiplex_network,3);
% 
% for i = 1:L
% 
%     one_out = multiplex_network(:,:,setdiff(1:L,i));
%     food_web = sum(one_out,3) > 0; 
% 
%     [trophic_web, trophic_species, trophic_species_indx, ~, isolated_species] ...
%         = unique_grouping(food_web);
% 
%     % remove isolated species for generating matching webs % this is NOT
%     % the taxonomic richness
%     trophic_web = trophic_web(~isolated_species,~isolated_species); 
% 
%     properties(1+i,:) = calculate_food_web_properties(trophic_web);
% 
% end
% 
% 
% % compare to the food web (all types of resources grouped together)
% food_web = double(logical(sum(multiplex_network,3) > 0)); 
% 
% [trophic_web, trophic_species, trophic_species_indx, ~, isolated_species] ...
%         = unique_grouping(food_web);
% 
%     % remove isolated species for generating matching webs % this is NOT
%     % the taxonomic richness
%     trophic_web = trophic_web(~isolated_species,~isolated_species); 
% 
%     properties(L+1,:) = calculate_food_web_properties(trophic_web);
% 
% 
% properties.Properties.RowNames = ["prey","flowers","leaves","seeds",...
%     "wood","carrion","food web"];
% 
%
% SCALE DEPENDENDE OF FOOD WEB STRUCTURE
% % compare the UMBS food web and other empirical food webs to simulated
% % niche webs as a null model to characterize their topologies 
% reps = 1000;
% 
% % get list of empirical webs 
% cd("empirical_trophic_webs/")
% file_directory = pwd; 
% cd ..
% 
% files = dir(fullfile(file_directory,'*.mat'));
% 
% for k = 1:length(files)
% 
%     % load each empirical web, stored as an edge list
%     base_name = files(k).name
%     full_name = fullfile(file_directory, base_name);
%     load(full_name, 'edge_list');
% 
%     S = max(edge_list,[],'all');
% 
%     % build food web from edge list
%     sparse_web = sparse(edge_list(:,1),edge_list(:,2),ones(length(edge_list),1),S,S);
%     food_web = full(sparse_web);
%     
%     % FOOD WEB AGGREGATED TO STRICT TROPHIC SPECIES
%     [trophic_web, trophic_species, trophic_species_indx, ~, isolated_species] ...
%         = unique_grouping(food_web);
% 
%     % remove isolated species for generating matching webs % this is NOT
%     % the taxonomic richness
%     trophic_web = trophic_web(~isolated_species,~isolated_species); 
%     
%     % COMPARE PROPERTIES OF EMPIRICAL FOOD WEB TO SIMULATED NICHE MODELS
%     % WITH MATCHED TROPHIC SPECIES RICHNESS AND CONNECTANCE
%     [empirical_properties, simulated_webs, simulated_properties, ...
%         simulated_generality, simulated_vulnerability ] = ...
%         matching_niche_models(trophic_web, reps); % check seed vals w/in fn
% 
%     save_name = ['trophic_properties2_', base_name];
% 
%     save(save_name,'empirical_properties','simulated_webs',...
%         'simulated_properties','trophic_web','simulated_generality',...
%         'simulated_vulnerability','-v7.3')
% 
%     clear simulated_webs
%     
% end
% 
% % rerun the script until done, if necessary 
% for k = 1:length(files) 
% 
%     % load each empirical web, stored as an edge list
%     base_name = files(k).name
% 
%     save_name = ['trophic_properties2_', base_name];
%     
%     load(save_name)
% 
%     successes = size(simulated_webs,2)
% 
%     if successes < reps % change this to a while loop if necessary
% 
%         % COMPARE PROPERTIES OF EMPIRICAL FOOD WEB TO SIMULATED NICHE MODELS
%         % WITH MATCHED TROPHIC SPECIES RICHNESS AND CONNECTANCE
%         [empirical_properties, simulated_webs_new, simulated_properties_new, ...
%             simulated_generality, simulated_vulnerability ] = ...
%             matching_niche_models(trophic_web, reps - successes);
%     
%         simulated_webs = [simulated_webs, simulated_webs_new];
%         simulated_properties = [simulated_properties; simulated_properties_new];
% 
%         successes = size(simulated_webs,2)
% 
%         save(save_name,'empirical_properties','simulated_webs',...
%            'simulated_properties','trophic_web','simulated_generality',...
%            'simulated_vulnerability','-v7.3')
% 
%     end
% end
% 
% 
% % % compare the UMBS food web to simulated niche webs with matched basal 
% % % species as a null model to characterize their topologies 
% reps = 1000; 
% 
%     % load each empirical web, stored as an edge list
%     base_name = 'temperate_forest_UMBS.mat'
%     full_name = fullfile(file_directory, base_name);
%     load(full_name, 'edge_list');
% 
%     S = max(edge_list,[],'all');
% 
%     % build food web from edge list
%     sparse_web = sparse(edge_list(:,1),edge_list(:,2),ones(length(edge_list),1),S,S);
%     food_web = full(sparse_web);
%     
%     % FOOD WEB AGGREGATED TO STRICT TROPHIC SPECIES
%     [trophic_web, trophic_species, trophic_species_indx, ~, isolated_species] ...
%         = unique_grouping(food_web);
% 
%     % remove isolated species for generating matching webs % this is NOT
%     % the taxonomic richness
%     trophic_web = trophic_web(~isolated_species,~isolated_species); 
%     
%     % COMPARE PROPERTIES OF EMPIRICAL FOOD WEB TO SIMULATED NICHE MODELS
%     % WITH MATCHED TROPHIC SPECIES RICHNESS, CONNECTANCE, AND BASAL SPECIES RICHNESS
%     [empirical_properties, simulated_webs, simulated_properties] = ...
%         matching_niche_models_basal(trophic_web, reps);
% 
%     save_name = ['trophic_properties_basal_trophic_random_', base_name];
% 
%     save(save_name,'empirical_properties','simulated_webs',...
%         'simulated_properties','trophic_web','-v7.3')
% 
% % these will take forever to simulate and probably will need to be looped over 
% % using the previous code chunk
% 
% 
%     clear simulated_webs


file_directory = pwd;
file_wildcard = "trophic_properties_*";

[NMEs_table, raw_errors_table, empirical_properties_table, ...
    simulated_properties] = calc_model_errors(file_directory, ...
    file_wildcard);

create_publication_results_figures()

[ks_results_table] = ...
    calculate_kolmogorov_smirnov_tests(file_directory, file_wildcard);

create_degree_distribution_figures()


function create_publication_results_figures() 

NMEs = readtable('NMEs_results.csv','ReadRowNames',true);
NMEs = sortrows(NMEs,'S','ascend');
NMEs('temperate_forest_no_basal_animals',:) = []; % just for visualization 
NMEs.group = [repmat(["historical"], height(NMEs)-2,1); "UMBS";  "UMBS_basal"]; % "flowers"; "leaves"; "seeds"; "wood"; 

unique_groups = unique(NMEs.group,'stable')

NMEs = renamevars(NMEs,{'fracStrictHerb','fracOmni','fracStrictHerbLinks', ...
    'avgGenCons','avgGenStrictHerb','GenSDStrictHerb','avgVulBasal','fracTrophicOmni','fracCann'},...
    {'TL2','Omniv','TL2Link','meanGen','meanGenTL2','GenSDTL2','meanVulBasal','TrophicOmniv','Cannib'});

vars = {'B','I','T','TL2','Omniv','TL2Link','meanTL','meanGen','meanGenTL2','GenSD','GenSDTL2',...
    'VulSD','meanVulBasal','VulSDBasal'};

colors = [0.6350 0.0780 0.1840]; 

figure
tiledlayout(2,7)
set(gcf,'color','w');
for i = 1:numel(vars)
    nexttile
    hold on
    
    patch([1 10000 10000 1],[-1 -1 1 1],'k','FaceAlpha',0.1,'EdgeAlpha',0.1) % max(NMEs.S) max(NMEs.S) 
    scatter(NMEs.S(NMEs.group == unique_groups(1)),NMEs.(vars{i})(NMEs.group == "historical"),... % unique_groups(1)
        50,'filled','MarkerFaceAlpha',0.5,'MarkerEdgeColor',[0 0.4470 0.7410]) % ,'ko')
    scatter(NMEs.S(NMEs.group == unique_groups(2)),NMEs.(vars{i})(NMEs.group == "UMBS"),... % unique_groups(2)
        70,'filled','r^')
    scatter(NMEs.S(NMEs.group == unique_groups(3)),NMEs.(vars{i})(NMEs.group == "UMBS_basal"),... % unique_groups(3)
        70,'^','MarkerEdgeColor',colors(1,:),'LineWidth',1)
    
    ylabel(vars{i})
    xlabel('S')
    set(gca,'FontSize',12);
    set(gca,'XScale','log')
    xlim([10,10000])
% grid on

end

vars = {'B','I','T','TL2','TrophicOmniv','Cannib',...
'maxTL','GenSD','VulSD','LinkSD','PathLen','MaxSim'}; % ClustCoeff % Loop 

figure
tiledlayout(2,6)
set(gcf,'color','w');
for i = 1:numel(vars)
    nexttile
    hold on
    
    patch([1 10000 10000 1],[-1 -1 1 1],'k','FaceAlpha',0.1,'EdgeAlpha',0.1) % max(NMEs.S) max(NMEs.S) 
    scatter(NMEs.S(NMEs.group == unique_groups(1)),NMEs.(vars{i})(NMEs.group == "historical"),...
        50,'filled','MarkerFaceAlpha',0.5,'MarkerEdgeColor',[0 0.4470 0.7410]) % ,'ko')
    scatter(NMEs.S(NMEs.group == unique_groups(2)),NMEs.(vars{i})(NMEs.group == "UMBS"),...
        70,'filled','r^')
    scatter(NMEs.S(NMEs.group == unique_groups(3)),NMEs.(vars{i})(NMEs.group == "UMBS_basal"),...
        70,'^','MarkerEdgeColor',colors(1,:),'LineWidth',1)
    
    ylabel(vars{i})
    xlabel('S')
    set(gca,'FontSize',12);
    set(gca,'XScale','log')
    xlim([10,10000])
% grid on

end

figure
set(gcf,'color','w');
hold on

patch([1 5000 5000 1],[0.01 0.01 1 1],'k','FaceAlpha',0.1,'EdgeAlpha',0.1) % max(NMEs.S) max(NMEs.S) 
scatter(NMEs.S(NMEs.group == unique_groups(1)),NMEs.mean_MEs(NMEs.group == "historical"),...
    50,'filled','MarkerFaceAlpha',0.5,'MarkerEdgeColor',[0 0.4470 0.7410]) % ,'ko')
scatter(NMEs.S(NMEs.group == unique_groups(2)),NMEs.mean_MEs(NMEs.group == "UMBS"),...
    70,'filled','r^')
scatter(NMEs.S(NMEs.group == unique_groups(3)),NMEs.mean_MEs(NMEs.group == "UMBS_basal"),...
    70,'^','MarkerEdgeColor',[0.6350 0.0780 0.1840],'LineWidth',1)
ylabel('mean |Normalized Model Error|')
xlabel('S')
set(gca,'FontSize',12);
set(gca,'XScale','log')
xlim([10,5000])
set(gca,'YScale','log')
ylim([0.3,100])

end

function create_degree_distribution_figures()

load('trophic_properties_basal_trophic_first_temperate_forest_UMBS.mat',...
    'simulated_generality','simulated_vulnerability','trophic_web')

simulated_generality_basal = simulated_generality;
simulated_vulnerability_basal = simulated_vulnerability;

basal_indx = 1:size(trophic_web,1); 

load('trophic_properties_temperate_forest_UMBS.mat',...
    'simulated_generality','simulated_vulnerability','trophic_web')

empirical_generality = sort(sum(trophic_web,2),'descend');
empirical_vulnerability = sort(sum(trophic_web,1)','descend'); 

trophic_indx = 1:size(trophic_web,1); 

figure
tiledlayout(2,1)
nexttile
    set(gcf,'color','w');
    for i = 1:1000
        hold on
        scatter(trophic_indx, sort(simulated_generality(i,:),'descend'),'ko','MarkerEdgeAlpha',0.5)
        scatter(basal_indx, sort(simulated_generality_basal(i,:),'descend'),'MarkerEdgeAlpha',0.5,'MarkerEdgeColor',[0 0.4470 0.7410])
    end
    hold on
    scatter(trophic_indx,empirical_generality,'r^','filled') %,[0.6350 0.0780 0.1840], 'MarkerEdgeColor',[0.6350 0.0780 0.1840])
    set(gca,'FontSize',12);
    set(gca,'YScale','log')
    ylabel('Species Generality (In-Degree)')
    xlabel('Species Index, Ordered by Decreasing Generality')
nexttile
    set(gcf,'color','w');
    for i = 1:1000
        hold on
        scatter(trophic_indx, sort(simulated_vulnerability(i,:),'descend'),'ko','MarkerEdgeAlpha',0.5)
        scatter(basal_indx, sort(simulated_vulnerability_basal(i,:),'descend'),'MarkerEdgeAlpha',0.5,'MarkerEdgeColor',[0 0.4470 0.7410])
    end
    hold on
    scatter(trophic_indx,empirical_vulnerability,'r^','filled') , % 'MarkerEdgeColor',[0.6350 0.0780 0.1840]) % [0.6350 0.0780 0.1840],
    
    ylabel('Species Vulnerability (Out-Degree)')
    xlabel('Species Index, Ordered by Decreasing Vulnerability')
    
    set(gca,'FontSize',12);
    set(gca,'YScale','log')

end

function [empirical_properties, simulated_webs, simulated_properties, ...
    simulated_generality, simulated_vulnerability ...
    ] = matching_niche_models(empirical_web, reps) 

k_S = sum(empirical_web,1) + (sum(empirical_web,2)'); 
isolated_species = k_S == 0;

empirical_web = empirical_web(~isolated_species, ~isolated_species);

empirical_properties = calculate_food_web_properties(empirical_web); 

S = empirical_properties.S;
C = empirical_properties.C;

successes = 0; 
tries = 0;

seed_S = S; %  + 4; % round(S * 1.1);
seed_C = C; % C * 1.2;

simulated_generality = zeros(reps,S); % 
simulated_vulnerability = zeros(reps,S); % 

for i = 1:reps % while successes < reps & tries < reps * 10 % 
    
    tries = tries + 1; 

    % ORIGINAL NICHE MODEL 
    [simulated_web,success] = niche_model_trophic_species(seed_S, seed_C, S, C);

    % FASTER VERSION (DOESN'T TEST FOR UNIQUE TROPHIC SPECIES) 
    % [simulated_web,success] = niche_model(S, C);

    if success

        successes = successes + 1;

        simulated_webs(successes).niche_model = simulated_web;
        [simulated_properties(successes,:), generality, ...
            vulnerability] = calculate_food_web_properties(simulated_web); 
        simulated_generality(successes,:) = generality; 
        simulated_vulnerability(successes,:) = vulnerability; 

    end
    
end

successes

if successes == 0
    simulated_webs = [];
    simulated_properties = [];
    % disp('no success')
end

end

function [empirical_properties, simulated_webs, simulated_properties, ...
    simulated_generality, simulated_vulnerability ...
    ] = matching_niche_models_basal(empirical_web, reps)

% remove isolated nodes
k_S = sum(empirical_web,1) + (sum(empirical_web,2)'); 
isolated_species = k_S == 0;
empirical_web = empirical_web(~isolated_species, ~isolated_species);

empirical_properties = calculate_food_web_properties(empirical_web); 

S = empirical_properties.S;
C = empirical_properties.C;
P = round(empirical_properties.S * empirical_properties.B);

successes = 0; 
tries = 0;

% seed_S = 2896;
% seed_C = 0.057;
% seed_P = 927;

seed_S = round(S * 1.1); % S + 4; % seed_S = round(S * 1.115); % S + 4;
seed_C = C * 1.2; % seed_C = C * 1.05;
seed_P = floor(round(P * 1.58)); % seed_P = floor(round(P * 1.49)*0.98); 

simulated_generality = zeros(reps,S); % 
simulated_vulnerability = zeros(reps,S); % 

for i = 1:reps % while successes < reps & tries < reps * 10 % 
    
    tries = tries + 1; 

    % % BASAL-MATCHED NICHE MODEL
    [simulated_web,success] = niche_model_trophic_species_basal(seed_S, seed_C, seed_P, S, C, P);

    if success

        successes = successes + 1;

        simulated_webs(successes).niche_model = simulated_web;
        [simulated_properties(successes,:), generality, ...
            vulnerability] = calculate_food_web_properties(simulated_web); 
        simulated_generality(successes,:) = generality;
        simulated_vulnerability(successes,:) = vulnerability; 
        % generality, vulnerability 
    end
    
end

successes

if successes == 0
    simulated_webs = [];
    simulated_properties = [];
    % disp('no success')
end

end

function [trophic_web,success] = niche_model_trophic_species(S, C, St, Ct)
% using Allesina et al. 2008 correction
% S: number of species
% C: directed connnectance of network: C = L/S^2, L: number of linnks
% traits: an Sx1 vector of values between 0 and 1
% checks: – realized num links is within 3% of target
%         – no species are completely disconnected
% NOT checking: - groups of species completely disconnected from basal
%                 species
%               - no redundant speces (all species are trophic species)

min_C = 0.97*Ct; % floor(0.97*L) % for links
max_C = 1.03*Ct; % ceil(1.03*L) % for links

min_S = St; % floor(0.99*S);
max_S = St; % ceil(1.01*S); 

% tries to achieve success at all the checks 
tries = 1000;
success = 0; 

while tries > 0

    tries = tries - 1;

    % generate and sort by niche values: n_sorted
    niche_values = sort(rand(S,1),'ascend');

    % start by assembling the full possible network    
    food_web = zeros(S,S);

    % niche interaction probabibilities:
    % assign r_i = n_i * B_i = size of feeding range
    %   B_i is drawn uniformly from (0,1) using the beta distribution with
    %     parameters (alpha,beta)
    %   beta is assigned so that E[X] == 1/(1+beta) == 2C, i.e. beta = 1/(2*C)-1
    %   C is the target connetance
    %   alpha is 1 for simplicity    
    alpha = ones(S,1);
    beta = (S-1)/(2*S*C) - 1; % correction from Allesina et al. 2008 (see SI) % ORIGINAL: 1/(2*C) - 1;

    E = betarnd(alpha,beta);
    feeding_ranges = niche_values .* E;
    
    % assign c_i = center of feeding range drawn uniformly from [r_i/2, n_i]
    % c = (traits - (r ./ 2)) .* rand(S,1) + (r ./ 2);
    feeding_centers = (min(niche_values, 1 - feeding_ranges./2) - (feeding_ranges./2)) .* rand(S,1) + (feeding_ranges./2);
           
    % assign the lowest niche value species (1) to feeding range = 0 so 
    % there is always at least one basal species in every web 
    feeding_ranges(1) = 0;
    
    % assign food_web(i,j) = {1 if n_j is an element of [c_i - r_i/2, c_i + r_i/2], 0 otherwise} 
    % niche model
    for i = 1:S
        low_i = feeding_centers(i) - feeding_ranges(i)/2;
        high_i = feeding_centers(i) + feeding_ranges(i)/2;
        food_web(i,niche_values >= low_i & niche_values <= high_i) = 1;
    end

    % strict trophic grouping of food web
    trophic_web = unique_grouping(food_web);

    % degree distribution of all species
    k_S = sum(trophic_web,2) + sum(trophic_web,1)';

    % realized richness
    r_S = sum(k_S > 0);

    % realized connectance
    r_C = sum(trophic_web,"all")/r_S^2;

    % realized basal richness
    r_P = sum(sum(trophic_web,2) == 0);

    if r_C >= min_C && r_C <= max_C 
    % correct connectance

        if r_S >= min_S && r_S <= max_S

            % if all(k_S)
            % everyone has a partner

                % successful network!
                success = 1;

                % disp(max_tries)
                tries = 0;

            % end  

        end
        
    end
    
end % while

end

function [trophic_web,success] = niche_model_trophic_species_basal(S, C, P, St, Ct, Pt)
% using Allesina et al. 2008 correction
% S: number of species
% C: directed connnectance of network: C = L/S^2, L: number of linnks
% traits: an Sx1 vector of values between 0 and 1
% checks: – realized num links is within 3% of target
%         – no species are completely disconnected
% NOT checking: - groups of species completely disconnected from basal
%                 species
%               - no redundant speces (all species are trophic species)

min_C = 0.97*Ct; % floor(0.97*L) % for links
max_C = 1.03*Ct; % ceil(1.03*L) % for links

min_S = St; % floor(0.99*St); % 
max_S = St; % ceil(1.01*St); % 

min_P = Pt; % floor(0.999*Pt);
max_P = Pt; % ceil(1.001*Pt); 

% tries to achieve success at all the checks 
tries = 1000;
success = 0; 

while tries > 0

    tries = tries - 1;

    % generate and sort by niche values: n_sorted
    niche_values = sort(rand(S,1),'ascend');

    % start by assembling the full possible network    
    food_web = zeros(S,S);

    % niche interaction probabibilities:
    % assign r_i = n_i * B_i = size of feeding range
    %   B_i is drawn uniformly from (0,1) using the beta distribution with
    %     parameters (alpha,beta)
    %   beta is assigned so that E[X] == 1/(1+beta) == 2C, i.e. beta = 1/(2*C)-1
    %   C is the target connetance
    %   alpha is 1 for simplicity    
    alpha = ones(S,1);
    beta = (S-1)/(2*S*C) - 1; % correction from Allesina et al. 2008 (see SI) % ORIGINAL: 1/(2*C) - 1;

    E = betarnd(alpha,beta);
    feeding_ranges = niche_values .* E;
    
    % assign c_i = center of feeding range drawn uniformly from [r_i/2, n_i]
    % c = (traits - (r ./ 2)) .* rand(S,1) + (r ./ 2);
    feeding_centers = (min(niche_values, 1 - feeding_ranges./2) - (feeding_ranges./2)) .* rand(S,1) + (feeding_ranges./2);
           
    % assign the lowest niche value species (1) to feeding range = 0 so 
    % there is alway at least one basal species in every web 
    feeding_ranges(1) = 0;

    % assign 1 less than the number of correct basal species to have
    % feeding range = 0 also 
    % feeding_ranges(1:P) = 0; % (randi(S,[P-1,1])) = 0; 
    feeding_ranges(1) = 0;
    feeding_ranges(randi(S,[P-1,1])) = 0; 
    % feeding_ranges(1:floor(0.98*P)) = 0; % UMBS3 
    
    % assign food_web(i,j) = {1 if n_j is an element of [c_i - r_i/2, c_i + r_i/2], 0 otherwise} 
    % niche model
    for i = 1:S
        low_i = feeding_centers(i) - feeding_ranges(i)/2;
        high_i = feeding_centers(i) + feeding_ranges(i)/2;
        food_web(i,niche_values >= low_i & niche_values <= high_i) = 1;
    end

    % strict trophic grouping of food web
    trophic_web = unique_grouping(food_web);

    % degree distribution of all species
    k_S = sum(trophic_web,2) + sum(trophic_web,1)';

    % realized richness
    r_S = sum(k_S > 0);

    % realized connectance
    r_C = sum(trophic_web,"all")/r_S^2;

    % realized basal richness
    r_P = sum(sum(trophic_web,2) == 0);

%     [r_S, max_S, r_P, max_P]
%     [r_C, min_C, max_C]

   if r_C >= min_C && r_C <= max_C 
    % correct connectance

        if r_S >= min_S && r_S <= max_S
        % correct richness

            if r_P >= min_P && r_P <= max_P
            % correct basal richness

                % successful network!
                success = 1;

                % disp(max_tries)
                tries = 0;

            end  

        end
        
    end
    
end % while

end

function [food_web,success] = niche_model(S, C) 
% using Allesina et al. 2008 correction
% S: number of species
% C: directed connnectance of network: C = L/S^2, L: number of linnks
% traits: an Sx1 vector of values between 0 and 1
% checks: – realized num links is within 3% of target
%         – no species are completely disconnected
% NOT checking: - groups of species completely disconnected from basal
%                 species
%               - no redundant speces (all species are trophic species)

min_C = 0.97*C; % floor(0.97*L) % for links
max_C = 1.03*C; % ceil(1.03*L) % for links

min_S = S; % floor(0.99*S);
max_S = S; % ceil(1.01*S); 

% tries to achieve success at all the checks 
tries = 1000;
success = 0; 

while tries > 0

    tries = tries - 1;

    % generate and sort by niche values: n_sorted
    niche_values = sort(rand(S,1),'ascend');

    % start by assembling the full possible network    
    food_web = zeros(S,S);

    % niche interaction probabibilities:
    % assign r_i = n_i * B_i = size of feeding range
    %   B_i is drawn uniformly from (0,1) using the beta distribution with
    %     parameters (alpha,beta)
    %   beta is assigned so that E[X] == 1/(1+beta) == 2C, i.e. beta = 1/(2*C)-1
    %   C is the target connetance
    %   alpha is 1 for simplicity    
    alpha = ones(S,1);
    beta = (S-1)/(2*S*C) - 1; % correction from Allesina et al. 2008 (see SI) % ORIGINAL: 1/(2*C) - 1;

    E = betarnd(alpha,beta);
    feeding_ranges = niche_values .* E;
    
    % assign c_i = center of feeding range drawn uniformly from [r_i/2, n_i]
    % c = (traits - (r ./ 2)) .* rand(S,1) + (r ./ 2);
    feeding_centers = (min(niche_values, 1 - feeding_ranges./2) - (feeding_ranges./2)) .* rand(S,1) + (feeding_ranges./2);
           
    % assign the lowest niche value species (1) to feeding range = 0 so 
    % there is always at least one basal species in every web 
    feeding_ranges(1) = 0;
    
    % assign food_web(i,j) = {1 if n_j is an element of [c_i - r_i/2, c_i + r_i/2], 0 otherwise} 
    % niche model
    for i = 1:S
        low_i = feeding_centers(i) - feeding_ranges(i)/2;
        high_i = feeding_centers(i) + feeding_ranges(i)/2;
        food_web(i,niche_values >= low_i & niche_values <= high_i) = 1;
    end

%     % strict trophic grouping of food web
%     trophic_web = unique_grouping(food_web);

    % degree distribution of all species
    k_S = sum(food_web,2) + sum(food_web,1)';

    % realized richness
    r_S = sum(k_S > 0);

    % realized connectance
    r_C = sum(food_web,"all")/r_S^2;

    % realized basal richness
    r_P = sum(sum(food_web,2) == 0);

    if r_C >= min_C && r_C <= max_C 
    % correct connectance

        if r_S >= min_S && r_S <= max_S

        % if all(k_S)
            % everyone has a partner

            % if size(trophic_web,1) == St ||  (size(trophic_web,1) == St+1) % size(trophic_web,1) == S || 
                % everyone is a unique trophic species

                % successful network!
                success = 1;

                % disp(max_tries)
                tries = 0;
           %  end
        % end  

        end
        
    end
    
end % while

if success == 0
    % disp('Unable to construct network in 1,000 tries.')
    food_web = zeros(S,S);
end

end

function [properties,generality,vulnerability] = calculate_food_web_properties(food_web)
% calls calc_TLs
% food web is of the form that row i eats column j

% species richness
S = size(food_web,1);
% number links
L = sum(food_web,'all');
% directed connectance
C = L/S^2; 
% link density
LD = sum(food_web,'all')/S; 

% logically index species 
plants = sum(food_web,2) == 0; % basal 
animals = ~plants; % consumers (intermediate & top) 
herbivores = sum(food_web(:,plants),2) > 0;
carnivores = sum(food_web(:,animals),2) > 0;
resources = (sum(food_web,1))' > 0; % (basal & intermediate) 
% prey = (sum(food_web(:,animals),1))' > 0;
top = (sum(food_web,1) == 0)' & animals; % imposing this extra constraint
isolated = (sum(food_web,1) == 0)' & plants;

% realized species richness
r_S = S - sum(isolated);

% calculate trophic levels [from the largest connected component]
[swTL, paTL, sTL] = calc_TLs(food_web); 

% calculate degree distributions of all species
generality = sum(food_web,2);
vulnerability =  sum(food_web,1)';

k_dist = generality + vulnerability; 

% SPECIES COMPOSITION
% fraction basal species (no prey)
B = sum(plants)/S;
% fraction top species (no predators, not isolated)
T = sum(top)/S; 
% fraction intermediate species
I = 1 - B - T; 
% fraction isolated species (no predators or prey) 
fracIsol = sum(isolated)/S; 

% fraction herbivore species
fracHerb = sum(herbivores)/S;
% fraction strict herbivore species, aka Herbiv
fracStrictHerb = sum(herbivores & ~carnivores)/S;
% fraction carnivore species 
fracCarn = sum(carnivores)/S;
% fraction strict carnivore species
fracStrictCarn = sum(~herbivores & carnivores)/S;
% fraction omnivore species (eat plants & animals)
fracOmni = sum(herbivores & carnivores)/S; 
% fraction cannibalistic species (eat themselves), aka Cannib
fracCann = sum(diag(food_web))/S;
% fraction trophic omnivore species, aka Omniv
% species with non-integer trophic levels & > 1 resource
integerTLs = rem(paTL,1) == 0; 
fracTrophicOmni = sum(generality > 1 & ~integerTLs)/S; 

% predator-prey ratio 
PPratio = (T+I)/(B+I);


% LINK COMPOSITION 
% fraction herbivory links
fracHerbLinks = sum(food_web(herbivores,plants),"all")/L;
% fraction strictherbivory links
fracStrictHerbLinks = sum(food_web(herbivores & ~carnivores,plants),"all")/L;
% fraction carnivory links
fracCarnLinks = sum(food_web(carnivores,animals),"all")/L;
% fraction carnivory links
fracStrictCarnLinks = sum(food_web(carnivores & ~herbivores,animals),"all")/L;


% LINK HETEROGENEITY (DEGREE DISTRIBUTION 
% average generality (diet breath) % should potentially also be normalized
% by LD? 
% mean number resource species for all species
avgGen = mean(generality); 
% mean number resource species for all consumers
avgGenCons = mean(generality(animals,:)); 
% mean number resource species for intermediate species 
avgGenInter = mean(generality(animals & ~top,:)); 
% mean number resource species for herbivores
avgGenHerb = mean(generality(herbivores,:)); 
% mean number resource species for strict herbivores 
avgGenStrictHerb = mean(generality(herbivores & ~carnivores,:)); 
% mean number resource species for carnivores
avgGenCarn = mean(generality(carnivores,:)); 
% mean number resource species for strict carnivores
avgGenStrictCarn = mean(generality(~herbivores & carnivores,:)); 


% heterogeneity in generality (diet breath) 
% % standard deviation of number resource species for all species
% rawGenSD = std(diet_breadth); 
% normalized by link density (LD) to control for web complexity
GenSD = std(generality)/LD; 
% std of number resource species for all consumers
GenSDCons = std(generality(animals,:)) / avgGenCons; 
% std of number resource species for intermediate species 
GenSDInter = std(generality(animals & ~top,:)) / avgGenInter; 
% std of number resource species for herbivores
GenSDHerb = std(generality(herbivores,:)) / avgGenHerb; 
% std of number resource species for strict herbivores 
GenSDStrictHerb = std(generality(herbivores & ~carnivores,:)) / avgGenStrictHerb; 
% std of number resource species for carnivores
GenSDCarn = std(generality(carnivores,:)) / avgGenCarn; 
% std of number resource species for strict carnivores
GenSDStrictCarn = std(generality(~herbivores & carnivores,:)) / avgGenStrictCarn; 

% average heterogeneity in trophic levels of diet, aka average
% "species omnivory": variance of trophic levels of a species' prey set
% % paTL_mat = repmat(paTL',[S,1]) .* food_web;
% % paTL_mat(paTL_mat == 0) = NaN; % omit nans (no feeding link)

% mean omnivory of consumers
% % avgOmni = mean(var(paTL_mat(animals,:), 0, 2,'omitnan'),'omitnan'); % for some reason this direct version doesn't work always? have to do it manually with a for loop 
avgOmni = calculate_average_omnivory(animals,food_web,paTL);
% mean omnivory of intermediate species 
avgOmniInter = calculate_average_omnivory(animals & ~top,food_web,paTL);
% mean omnivory of herbivores
avgOmniHerb = calculate_average_omnivory(herbivores,food_web,paTL);
% mean omnivory of strict herbivores % should be 0 % calculating as a test
avgOmniStrictHerb = calculate_average_omnivory(herbivores & ~carnivores,food_web,paTL);
if avgOmniStrictHerb ~= 0 && ~isnan(avgOmniStrictHerb)
    disp('pause')
end
% mean omnivory of carnivores
avgOmniCarn = calculate_average_omnivory(carnivores,food_web,paTL);
% mean omnivory of strict carnivores
avgOmniStrictCarn = calculate_average_omnivory(carnivores & ~herbivores,food_web,paTL);


% average vulnerability (predator richness)
% mean number consumer species for all species % should potentially also be
% normalized by LD? 
avgVul = mean(vulnerability); 
if avgGen ~= LD
    disp('error')
end
if avgVul ~= LD
    disp('error')
end
% mean number consumer species for all resources
avgVulRes = mean(vulnerability(resources,:)); 
% mean number consumer species for basal species
avgVulBasal = mean(vulnerability(plants,:)); 
% mean number consumer species for intermediate species 
avgVulInter = mean(vulnerability(animals & ~top,:)); 
% mean number consumer species for herbivores 
avgVulHerb = mean(vulnerability(herbivores,:));
% mean number consumer species for strict herbivores 
avgVulStrictHerb = mean(vulnerability(herbivores & ~carnivores,:)); 
% basal out-degree (herbivory) link density
LDh = sum(food_web(herbivores,plants),"all")/sum(plants); 
if avgVulBasal ~= LDh
    disp('pause')
end


% heterogeneity in vulnerability (predator richness) 
% % standard deviation of number consumer species for all species
% rawVulSD = std(vulnerability); 
% normalized by link density (LD) to control for web complexity
VulSD = std(vulnerability)/LD; 
% std of number consumer species for all resources
VulSDRes = std(vulnerability(resources,:)) / avgVulRes; % LD of consumers per all resources
% std of number consumer species for basal species
VulSDBasal = std(vulnerability(plants,:)) / avgVulBasal; % LD of consumers per all plants
% std of number consumers species for intermediate species 
VulSDInter = std(vulnerability(animals & ~top,:)) / avgVulInter; % LD of consumers per all intermediate species
% std of number consumer species for herbivores
VulSDHerb = std(vulnerability(herbivores,:)) / avgVulHerb; % LD of consumers per all herbivores
% std of number consumer species for strict herbivores 
VulSDStrictHerb = std(vulnerability(herbivores & ~carnivores,:)) / avgVulStrictHerb; % LD of consumers per all strict herbivores


% heterogeneity in all interactions (consumer + resource richness)
% % standard deviation of total degree for all species
% rawLinkSD = std(k_dist); 
% normalized by link density (LD) to control for web complexity
LinkSD = std(k_dist)/LD; 


% HIERARCHY
% mean trophic level of consumers
meanTL = mean(swTL(animals),'omitnan'); % mean(paTL(animals));
% standard deviation of trophic level among consumers
SDTL = std(swTL(animals),'omitnan');
% max trophic level of consumers
maxTL = max(swTL); 
% mean trophic level of top consumers
meanTLtop = mean(swTL(top),'omitnan'); % mean(paTL(top_sp));

% minimum path length between each basal species and each other species 
% excludes cannibalism, loops, and infs where there is no path
food_web_graph = digraph(food_web'); % my graph flows are pointing the reverse direction
d = distances(food_web_graph,find(plants)); 

% characteristic path length, average of minimum directed 
% path lengths, aka PathLen
PathLen = mean(d(~isinf(d)));
% standard deviation of path lengths
PathSD = std(d(~isinf(d)));
% log of total directed paths beween basal and other species
logNumPaths = log(sum(~isinf(d),'all'));

% % clustering coefficient
% triangles = allcycles(food_web_graph,'MaxCycleLength',3,'MinCycleLength',3);
% triangle_mat = unique(cell2mat(triangles),'rows');
% 
% if max(size(triangle_mat)) ~= max(size(triangles))
%     disp('pause')
% end
% 
% denom = (S * (S-1) * (S-2))/(3 * 2); 
% ClustCoeff = size(triangle_mat,1)/denom;

% NICHE PARTITIONING 
% mean maximum trophic similarity
D = pdist(food_web,'jaccard'); % trophic similarity between each pair of species
D_matrix = squareform(D);

max_species_trophic_similarity = 1 - min(D_matrix + eye(S),[],2,'omitnan');
MaxSim = mean(max_species_trophic_similarity);

% aggregated to strict trophic species
trophic_web = unique_grouping(food_web);

% trophic species richness
t_S = size(trophic_web,1); 

% trophic species web connectance 
t_C = sum(trophic_web,'all')/(t_S*t_S);

% % entropy: measures the diversity of pathways of energy flow between 
% % species in the network
% rooted_food_web = zeros(S+1, S+1); % let basal species eat a hypothetical 'root'
% rooted_food_web(1:S,1:S) = food_web;
% rooted_food_web(plants, S+1) = 1; 
% 
% lambda = eigs(rooted_food_web,1); % find dominant eigenvalue
% H = log(lambda); % natural log
% 
% % scaled entropy: accounting for increase in nodes
% Hscaled = H/log(S+1); 


properties = table(S, C, L, LD, r_S, t_S, t_C, ...
    B, I, T, PPratio, ... 
    fracHerb, fracStrictHerb, fracCarn, fracStrictCarn, ...
    fracOmni, fracTrophicOmni, fracCann, ...
    fracHerbLinks, fracStrictHerbLinks, fracCarnLinks, fracStrictCarnLinks, ... % avgGen % avgVul 
    avgGenCons, avgGenInter, avgGenHerb, ...
    avgGenStrictHerb, avgGenCarn, avgGenStrictCarn, ...
    GenSD, GenSDCons, GenSDInter, GenSDHerb, ...
    GenSDStrictHerb, GenSDCarn, GenSDStrictCarn, ...
    avgOmni, avgOmniInter, avgOmniHerb, avgOmniCarn, ...
    avgOmniStrictCarn, ...
    avgVulRes, avgVulBasal, avgVulInter, avgVulHerb, ...
    avgVulStrictHerb,...
    VulSD, VulSDRes, VulSDBasal, VulSDInter, ...
    VulSDHerb, VulSDStrictHerb, ...
    LinkSD, ...
    meanTL, SDTL, maxTL, meanTLtop, PathLen, PathSD, ...
    logNumPaths, ...
    MaxSim); % , ClustCoeff); 

generality = generality(k_dist>0)';
vulnerability = vulnerability(k_dist>0)';

end

function [avgOmni] = calculate_average_omnivory(logical_index,food_web,paTL)

indx = find(logical_index);

var_paTL = zeros(numel(indx),1);

for i = 1:numel(indx)
    var_paTL(i,1) = var(paTL(logical(food_web(indx(i),:))));
end

avgOmni = mean(var_paTL,'omitnan');

end

function [NMEs_results, raw_error_results, empirical_properties_table, ...
    simulated_properties_table] = calc_model_errors(file_directory, file_wildcard)

files = dir(fullfile(file_directory,file_wildcard)); 

load(fullfile(file_directory, files(1).name), 'empirical_properties')

num_files = length(files);
num_properties = width(empirical_properties);

% normalized model errors % within +/-1 is significantly different
% normalized by the difference between the model median and 97.5 or 2.5
% percentile
NMEs = zeros(num_files,num_properties);
raw_errors = zeros(num_files,num_properties); 
mean_simulated_properties = zeros(num_files,num_properties); 

for k = 1:length(files)

    base_name = files(k).name
    full_name = fullfile(file_directory, base_name);
    
    web_names(k,1) = string(extractAfter(extractBefore(files(k).name,'.mat'),'trophic_properties_'));

    load(full_name, 'simulated_properties', ... % 'trophic_web', 'simulated_webs',
        'empirical_properties')

    empirical_properties_table(k,:) = empirical_properties;

    for i = 1:width(simulated_properties)

        % Normalized Model Errors (NMEs)
        % calculate 2.5 & 97.5 percentile and median of simulated data 
        % (50 percentile)
        simulated_percentiles = prctile(simulated_properties.(i), ...
            [2.5 50 97.5]);
        empirical_value = empirical_properties.(i);

        % compare empirical to 97.5 percentile if empirical val is larger 
        % than median of simulated
        if empirical_value >= simulated_percentiles(2)

            % use the positive sign to indicate that the true property is 
            % GREATER than the model estimation
            NMEs(k,i) = 1 * (simulated_percentiles(2) - empirical_value) / ...
                (simulated_percentiles(2) - simulated_percentiles(3));

        else % empirical_value < simulated_percentiles(2)

            % compare empirical to 2.5 percentile if empirical val is less 
            % than median of simulated
            % use the positive sign to indicate that the true property is 
            % LESS than the model estimation
            NMEs(k,i) = -1 * (simulated_percentiles(2) - empirical_value) / ...
                (simulated_percentiles(2) - simulated_percentiles(1));
        end
        
        NMEs(k,1) = empirical_properties.S; 
        NMEs(k,2) = empirical_properties.C; 

        % Raw Errors
        % calculate difference between mean of simulated data and empirical 
        % value
        raw_errors(k,i) = (mean(simulated_properties.(i),'omitnan') - ...
            empirical_value);
        mean_simulated_properties(k,i) = mean(simulated_properties.(i),'omitnan');

    end
end

NMEs_results = table(NMEs);
NMEs_results = splitvars(NMEs_results,'NMEs','NewVariableNames',simulated_properties.Properties.VariableNames);
NMEs_results.Properties.RowNames = web_names;
writetable(NMEs_results,'NMEs_results.csv',"WriteRowNames",true);

raw_error_results = table(raw_errors);
raw_error_results = splitvars(raw_error_results,'raw_errors','NewVariableNames',simulated_properties.Properties.VariableNames);
raw_error_results.Properties.RowNames = web_names; 
writetable(raw_error_results,'raw_error_results.csv',"WriteRowNames",true)

empirical_properties_table.Properties.RowNames = web_names;
writetable(empirical_properties_table,'empirical_properties.csv',"WriteRowNames",true)

simulated_properties_table = table(mean_simulated_properties); 
simulated_properties_table = splitvars(simulated_properties_table,'mean_simulated_properties','NewVariableNames',simulated_properties.Properties.VariableNames);
simulated_properties_table.Properties.RowNames = web_names;
writetable(simulated_properties_table,'simulated_properties_table.csv',"WriteRowNames",true)

NMEs_results.B(isnan(NMEs_results.B)) = 0;
NMEs_results.T(isnan(NMEs_results.T)) = 0;

mean_MEs = (abs(NMEs_results.T) + abs(NMEs_results.I) + abs(NMEs_results.B) ...
    + abs(NMEs_results.fracCann) + abs(NMEs_results.fracStrictHerb) ...
    + abs(NMEs_results.fracTrophicOmni) + abs(NMEs_results.LinkSD) ...
    + abs(NMEs_results.GenSD) + abs(NMEs_results.VulSD) ...
    + abs(NMEs_results.PathLen) + abs(NMEs_results.meanTL) ...
    + abs(NMEs_results.MaxSim)) / 12; % excluding loop due to how big our web is...
% missing ClustCoeff (Clustering Coefficient), Loop (Fraction Species in Loops)

NMEs_results.mean_MEs = mean_MEs; 
writetable(NMEs_results,'NMEs_results.csv',"WriteRowNames",true);

end

function [KS_test_results] = calculate_kolmogorov_smirnov_tests(file_directory, file_wildcard)

load('trophic_properties_temperate_forest_UMBS.mat','trophic_web')

[trophic_web,~,~,~,isolated_logical] = unique_grouping(trophic_web);
trophic_web = trophic_web(~isolated_logical,~isolated_logical); 

files = dir(fullfile(file_directory,file_wildcard)); 
num_files = length(files);

% compare each web's degree distributions to the UMBS webs degree
% distributions using two-sample Kolmogorov-Smirnov tests
UMBS_generality = sum(trophic_web,2); 
UMBS_vulnerability = sum(trophic_web,1)'; 

% 'sample size' of the UMBS web (number of species, i.e., degrees in the
% distribution) 
N1 = length(UMBS_generality); 

% testing the null hypothesis that the degree distribution was sampled from
% the same population as another sample, potentially of a different size
H_UMBS_generality = zeros(num_files,1);
H_UMBS_vulnerability = zeros(num_files,1);

% the null hypothesis will be rejected at the 0.05 significance level, for a
% two-tailed test; that is at 0.025 for a lower or higher cumulative
% probability 
P_UMBS_generality = zeros(num_files,1); 
P_UMBS_vulnerability = zeros(num_files,1); 

% the asymptotic P-value becomes very accurate for large sample sizes, and
% is believed to be reasonably accurate for sample sizes N1 and N2 such 
% that (N1*N2)/(N1 + N2) >= 4.
ks_sample_UMBS = zeros(num_files,1);

% also compare each empirical web's degree distributions to its simulated degree 
% distributions using two-sample Kolmogorov-Smirnov tests

% fraction of times the null hypothesis was rejected at the 0.05
% significance level in the ensemble of simulated webs
frac_rejected_generality = zeros(num_files,1);
frac_rejected_vulnerability = zeros(num_files,1); 

mean_Ps_generality = zeros(num_files,1);
mean_Ps_vulnerability = zeros(num_files,1); 

% also check if the asymptotic P-values are accurate, but here it will be
% (N2*N2)/(N2 + N2) >= 4 since the simulated distributions are the same
% size as the empirical web
ks_sample = zeros(num_files,1); 

% load in each empirical web
for k = 1:length(files)

    base_name = files(k).name
    full_name = fullfile(file_directory, base_name);
    
    web_names(k,1) = string(extractAfter(extractBefore(files(k).name,'.mat'),'trophic_properties_'));

    % and their simulated in- and out-degree distributions (aka generality
    % and vulnerability, respectively) 
    load(full_name, 'trophic_web', 'simulated_generality', 'simulated_vulnerability')

    % format the empirical webs & degree distributions
    [trophic_web,~,~,~,isolated_logical] = unique_grouping(trophic_web);
    trophic_web = trophic_web(~isolated_logical,~isolated_logical); 

    empirical_generality = sum(trophic_web,2); 
    empirical_vulnerability = sum(trophic_web,1)'; 

    % compare to the UMBS degree distributions
    [H_UMBS_generality(k,1),P_UMBS_generality(k,1)] = ...
        kstest2(UMBS_generality,empirical_generality);
    [H_UMBS_vulnerability(k,1),P_UMBS_vulnerability(k,1)] = ...
        kstest2(UMBS_vulnerability,empirical_vulnerability); 

    N2 = length(empirical_generality);
    ks_sample_UMBS(k) = (N1*N2)/(N1 + N2); % >= 4;

    % compare to the ensemble of simulated degree distributions 
    ks_sample(k,1) = (N2 * N2)/(N2 + N2); % >= 4;

    % num simulations
    reps = size(simulated_generality,1);

    Hs_generality = zeros(reps,1); Ps_generality = zeros(reps,1); 
    Hs_vulnerability = zeros(reps,1); Ps_vulnerability = zeros(reps,1); 

    for i = 1:reps

        [Hs_generality(i),Ps_generality(i)] = kstest2(empirical_generality,simulated_generality(i,:)); 
        [Hs_vulnerability(i),Ps_vulnerability(i)] = kstest2(empirical_vulnerability,simulated_vulnerability(i,:)); 
    
    end

    frac_rejected_generality(k,1) = sum(Hs_generality)./reps;
    frac_rejected_vulnerability(k,1) = sum(Hs_vulnerability)./reps; 
    mean_Ps_generality(k,1) = sum(Ps_generality)./reps; 
    mean_Ps_vulnerability(k,1) = sum(Ps_vulnerability)./reps; 

end

KS_test_results = table(H_UMBS_generality,P_UMBS_generality,...
    H_UMBS_vulnerability,P_UMBS_vulnerability,ks_sample_UMBS,...
    frac_rejected_generality,mean_Ps_generality,...
    frac_rejected_vulnerability,mean_Ps_vulnerability,ks_sample);
KS_test_results.Properties.RowNames = web_names;
writetable(KS_test_results,'KS_test_results.csv',"WriteRowNames",true);

end
```
