## Supplementary material for "A highly resolved network reveals the role of terrestrial herbivory in structuring aboveground food webs": Data and Code: taxonomic_web_analysis.m

```
% load species list from file
species_list = readtable('species_list_EDI.csv','TextType','String');

% load interaction list from file
data = readtable('interaction_list_EDI.csv','TextType','String'); 

interaction_types = unique(data.Interaction)
% this should match the resource that are plant vs animal types
% exclusively, to remove interactions that are ambiguous due to duplicated
% genus names used across taxonomic groups
plant_resource_indx = logical([0 1 1 1 1 0]');
num_resources = length(interaction_types);

% look at species composition
groupcounts(species_list,{'groups','orders'}) % use 'groups', 'families', 'genera' etc to see further breakdown

% fraction of 'species' that are recorded to genus level (ln 157)
fraction_genera = sum(strcmp(species_list.genera, species_list.species_names))/height(species_list);

% % stats on species composition
% groupcounts(species_list,{'groups'}) 
num_orders = height(unique(species_list.orders));
num_families = height(unique(species_list.families));
num_genera = height(unique(species_list.genera));

% assign species indices and record the indices of the consumer & resource
% species for each entry in the interaction list
S = height(species_list) % richness of the food web
species_list.indx = (1:S)';

num_records = height(data) % number of records in the interaction list (ln 309)

% index the interaction list % [function] 
data = index_interaction_list(data,species_list);

% record the number of links in the food web generated by each entry in the
% interaction list
data.NumLinks = cellfun(@numel, data.ConsumerIndx) .* cellfun(@numel, data.ResourceIndx);

% build and analyze the multiplex food web % [function] 
[species_list, richnesses, linkages, ...
    resource_type_usage, multifeeding, expanded_edge_list, ...
    multiplex_network] = analyze_edge_list(data, species_list, plant_resource_indx);

% resource_types = {"prey","flowers","leaves","seeds","wood","carrion"}';
% edge_list = table(species_list.species_names(expanded_edge_list(:,1)),...
%     species_list.species_names(expanded_edge_list(:,3)),...
%     resource_types(expanded_edge_list(:,2)),...
% 'VariableNames',{'ConsumerSpecies','ResourceSpecies','ResourceType'});
% writetable(edge_list,'edge_list_EDI.csv')


% % visualize diet breadths on different resources
% create_diet_breadth_plots(multiplex_network, species_list, data)

% look at species without interaction records
in_degree = sum(sum(multiplex_network,3),2); % total num resources each species has
out_degree = sum(sum(multiplex_network,3),1); % total num consumers each species has

degree_distribution = in_degree + out_degree'; % total num interactions each species has

species_list.SumInDegree = in_degree;
species_list.SumOutDegree = out_degree';

% species with no records at all 
species_with_no_records = species_list(degree_distribution == 0,:); % no interactions
groupsummary(species_with_no_records,{'groups','orders','families','genera'})
num_species_with_no_records = height(species_with_no_records)

% no records of resources
animals_with_no_resources = species_list(in_degree == 0 ... % no resources
    & ~(species_list.groups == "Plants"),:); % animal
groupsummary(animals_with_no_resources,{'groups','orders'});
num_animals_with_no_resources = height(animals_with_no_resources)

% no records that reference them by binomial name 
recorded_taxa = unique([data.Consumer; data.Resource]);
[~,indx] = setdiff(species_list.species_names,recorded_taxa);
species_with_no_specific_records = species_list(indx,:);
groupsummary(species_with_no_specific_records,{'groups','orders'});
num_species_with_no_specific_records = height(species_with_no_specific_records)

% records that reference them by binomial name 
all_species = sortrows(species_list,'species_names','ascend');
species_level = ismember(all_species.species_names,unique([data.Consumer; data.Resource]));
frac_species_referenced_as_binomials = sum(species_level)/S

all_species = all_species(~species_level,:);

% records that reference them as genera
genus_level = ismember(all_species.genera,unique([data.Consumer; data.Resource]));
frac_species_referenced_as_genus = sum(genus_level)/S

all_species = all_species(~genus_level,:)

% records that reference them as families
family_level = ismember(all_species.families,unique([data.Consumer; data.Resource]));
frac_species_referenced_as_family = sum(family_level)/S

all_species = all_species(~family_level,:)

% records that reference them as orders
order_level = ismember(all_species.orders,unique([data.Consumer; data.Resource]));
frac_species_referenced_as_order = sum(order_level)/S


% calculate connectance of each subnetwork using richnesses & linkages
% calculation of connectance depends on interaction type
linkages.Cx = zeros(num_resources+2,1);
linkages.Cx(~plant_resource_indx) = linkages.Lx(~plant_resource_indx)./(richnesses.Sx(~plant_resource_indx).^2); % carnivory
linkages.Cx(plant_resource_indx) = linkages.Lx(plant_resource_indx) ./ (richnesses.Ax(plant_resource_indx) .* richnesses.Px(plant_resource_indx)); % herbivory 

% expected food web connectance from a scalng law fit to empirical webs 
% Cohen & Briand 1984, Montoya & Solé 2003, Ings et al. 2009
C = 0.8 * richnesses.Sx(1)^(-0.5);

% look at the species that eat each resource
logical_consumers = species_list.InDegree > 0; 
num_resources_used = sum(logical_consumers,2);

frac_herbivores = sum(sum(species_list.InDegree(:,plant_resource_indx),2)>0)/richnesses.Conx("flattened_all"); % (ln 346)
frac_plant_resource_specialists = sum(resource_type_usage.NumConsumers(plant_resource_indx))/richnesses.Conx("flattened_all"); % (ln 347)
frac_resource_type_generalists = sum(resource_type_usage.NumConsumers(num_resources+1:end))/richnesses.Conx("flattened_all"); % (ln 360)
frac_carnivores = sum(sum(species_list.InDegree(:,~plant_resource_indx),2)>0)/richnesses.Conx("flattened_all"); % (ln 346)

% summary of species that eat animal tissues only (prey & carrion)
% remove semicolon to see the summary 
groupsummary(species_list((logical_consumers(:,1) | logical_consumers(:,6)) & num_resources_used <= 2,:),{'groups','orders'});  % (ln 348-9)
frac_carnivores = height(ans)/richnesses.Conx(end)

% summary of species that eat prey 
groupsummary(species_list(logical_consumers(:,1),:),{'groups','orders'});

% summary of species that eat carrion 
groupsummary(species_list(logical_consumers(:,6),:),{'groups','orders'});

% summary of species that eat flowers & leaves
groupsummary(species_list(logical_consumers(:,2) & logical_consumers(:,3),:),{'groups','orders'}); % (ln 361)

% summary of species that eat > 2 resource types
groupsummary(species_list(num_resources_used > 2,:),{'groups','orders'}); % (ln 362)

% look at relationships between feeding on multiple resource types
create_degree_correlation_plots(species_list,interaction_types);

% fraction of links where a consumer species is eating multiple types of the 
% same resource species % (ln 373)
frac_multifeeding_links = (linkages.Lx("multiplex") - linkages.Lx("flattened_binary"))/linkages.Lx("multiplex");

% summary of species that eat multiple tissues of the same plant
[consumers, resources] = find(sum(multiplex_network(:,:,plant_resource_indx),3) > 1);
groupsummary(species_list(unique(consumers),:), {'groups','orders'});

% look at the plants that supply these resources 
logical_resources = species_list.OutDegree > 0; 
num_resources_hosted = sum(logical_resources,2);

frac_multiresource_plants = sum(resource_type_usage.NumResources(num_resources+1:end))/richnesses.Px("flattened_all"); % (ln 374)

groupsummary(species_list(num_resources_hosted == 4,:),{'groups','orders'}); % (ln 362)

% look at trophic species groups - these are groups of species with the
% same consumers and resources
% first look at the ones only differentiated by species (traditional food
% web)
[trophic_food_web, trophic_species, trophic_food_web_indx, ~, isolated_logical] ...
    = unique_grouping(double(logical(sum(multiplex_network,3))));

S = size(trophic_food_web,1) - sum(isolated_logical) % this the trophic group of species with no links

species_list.trophic_species_indx = trophic_food_web_indx;

unique_trophic_web_species = species_list( ...
    ismember(species_list.trophic_species_indx, ...
    find(groupcounts(trophic_food_web_indx) == 1)),:); 

% now look at the ones also differentiated by resource type
[trophic_network, trophic_species, trophic_species_indx, ~, isolated_species] ...
    = unique_grouping(multiplex_network);

species_list.trophic_species_indx = trophic_species_indx;

unique_trophic_species = species_list( ...
    ismember(species_list.trophic_species_indx, ...
    find(groupcounts(species_list.trophic_species_indx) == 1)),:); 

num_trophic_species_diff_by_resource_type = length(trophic_network) -  length(trophic_food_web)

num_unique_trophic_species = height(unique_trophic_species)
height(unique_trophic_web_species);

groupsummary(unique_trophic_species,'groups');

species_in_trophic_groups = species_list( ...
    ismember(species_list.trophic_species_indx, ...
    find(groupcounts(species_list.trophic_species_indx) > 1)),:); 

species_in_big_trophic_groups = species_list( ...
    ismember(species_list.trophic_species_indx, ...
    find(groupcounts(species_list.trophic_species_indx) > 10)),:); 

num_species_in_big_trophic_groups = height(species_in_big_trophic_groups); 

sortrows(groupsummary(species_in_big_trophic_groups,{'trophic_species_indx','orders'}),'GroupCount','descend');

num_animals_with_no_resources_in_big_trophic_groups = sum(ismember(animals_with_no_resources.indx, species_in_big_trophic_groups.indx));
frac_animals_with_no_resources_in_big_trophic_groups = num_animals_with_no_resources_in_big_trophic_groups/num_animals_with_no_resources

frac_animals_in_big_trophic_groups_with_no_resources = numel(intersect(animals_with_no_resources.species_names, species_in_big_trophic_groups.species_names))/height(animals_with_no_resources)

% compare carnivory & herbivory subnetwork structure between original and
% trophic network
in_degree = sum(sum(multiplex_network(:,:,1),3),2);
out_degree = sum(sum(multiplex_network(:,:,1),3),1)';

in_degree_trophic = sum(sum(trophic_network(:,:,1),3),2);
out_degree_trophic = sum(sum(trophic_network(:,:,1),3),1)';

relative_carnivory_richness =  sum(in_degree_trophic > 0 | out_degree_trophic > 0) / sum(in_degree > 0 | out_degree > 0);
relative_carnivory_linkage = sum(in_degree_trophic) / sum(in_degree);

in_degree = sum(sum(multiplex_network(:,:,2:5),3),2);
out_degree = sum(sum(multiplex_network(:,:,2:5),3),1)';

in_degree_trophic = sum(sum(trophic_network(:,:,2:5),3),2);
out_degree_trophic = sum(sum(trophic_network(:,:,2:5),3),1)';

relative_herbivory_richness = sum(in_degree_trophic > 0 | out_degree_trophic > 0) / sum(in_degree > 0 | out_degree > 0);
relative_herbivory_linkage = sum(in_degree_trophic) / sum(in_degree);

% look at the level of taxonomic resolution different interactions are
% being sourced from 
check_edge_source(data,species_list,plant_resource_indx)

function [indexed_interaction_list] = ...
    index_interaction_list(interaction_list, species_list)

indexed_interaction_list = interaction_list; 

indexed_interaction_list.ConsumerIndx(interaction_list.ConsumerLevel == "Order") = ...
    arrayfun(@(x) species_list.indx(strcmpi(x, species_list.orders)), ...
   interaction_list.Consumer(interaction_list.ConsumerLevel == "Order"), 'UniformOutput', false);
indexed_interaction_list.ResourceIndx(interaction_list.ResourceLevel == "Order") = arrayfun(@(x) species_list.indx(strcmpi(x, species_list.orders)), interaction_list.Resource(interaction_list.ResourceLevel == "Order"), 'UniformOutput', false);

indexed_interaction_list.ConsumerIndx(interaction_list.ConsumerLevel == "Family") = ...
    arrayfun(@(x) species_list.indx(strcmpi(x, species_list.families)), ...
    interaction_list.Consumer(interaction_list.ConsumerLevel == "Family"), 'UniformOutput', false);
indexed_interaction_list.ResourceIndx(interaction_list.ResourceLevel == "Family") = arrayfun(@(x) species_list.indx(strcmpi(x, species_list.families)), ...
    interaction_list.Resource(interaction_list.ResourceLevel == "Family"), 'UniformOutput', false);

indexed_interaction_list.ConsumerIndx(interaction_list.ConsumerLevel == "Genus") = ...
    arrayfun(@(x) species_list.indx(strcmpi(x, species_list.genera)), ...
    interaction_list.Consumer(interaction_list.ConsumerLevel == "Genus"), 'UniformOutput', false);
indexed_interaction_list.ResourceIndx(interaction_list.ResourceLevel == "Genus") = ...
    arrayfun(@(x) species_list.indx(strcmpi(x, species_list.genera)), ...
    interaction_list.Resource(interaction_list.ResourceLevel == "Genus"), 'UniformOutput', false);

indexed_interaction_list.ConsumerIndx(interaction_list.ConsumerLevel == "Species") = ...
    arrayfun(@(x) species_list.indx(strcmpi(x, species_list.species_names)), ...
    interaction_list.Consumer(interaction_list.ConsumerLevel == "Species"), 'UniformOutput', false);
indexed_interaction_list.ResourceIndx(interaction_list.ResourceLevel == "Species") = ...
    arrayfun(@(x) species_list.indx(strcmpi(x, species_list.species_names)), ...
    interaction_list.Resource(interaction_list.ResourceLevel == "Species"), 'UniformOutput', false);


end

function [species_list_degs, richnesses, linkages, ...
    resource_type_usage, multifeeding_links, expanded_edge_list, ...
    multiplex_network] = analyze_edge_list(edge_list, species_list, plant_resource_indx)

% % Assemble Multiplex Food Web from Interaction List
interaction_types = unique(edge_list.Interaction);
N = size(interaction_types,1); % num interaction types
interaction_indx = (1:N)';
edge_list.InteractionIndx = arrayfun(@(x) interaction_indx(strcmpi(x, interaction_types)), ...
    edge_list.Interaction, 'UniformOutput', false);

% create an edge list of links between consumer species indices and resource
% species indices
temp_consumer = edge_list.ConsumerIndx;
temp_interactions = edge_list.InteractionIndx;
temp_resources = edge_list.ResourceIndx;

% some interaction records involve multiple species (e.g., all species in a
% genus)
% expand edges to all resource species
expanded_resources = cell2mat(temp_resources);
num_resources = cellfun(@length, temp_resources);
expanded_consumer = repelem(temp_consumer, num_resources); % species: the (species) indx of the original partner set % the second argument specifies how many times to repeat each item in the first array
expanded_interactions = repelem(temp_interactions, num_resources); 

% expand edges to all consumer species
temp_consumer = expanded_consumer;
temp_interactions = expanded_interactions;
temp_resources = expanded_resources;

expanded_consumer = cell2mat(temp_consumer);
num_focal = cellfun(@length, temp_consumer);
expanded_resources = repelem(temp_resources, num_focal); % species: the (species) indx of the original partner set % the second argument specifies how many times to repeat each item in the first array
expanded_interactions = cell2mat(repelem(temp_interactions, num_focal)); 

% this is the final edge list
expanded_edge_list = unique([expanded_consumer, expanded_interactions, expanded_resources],'rows');

% assemble a subnetwork for each interaction type from the final edge list 
% calculate stats on each subnetwork at the same time
species_list = sortrows(species_list,'indx','ascend');
S = height(species_list); % number of species (max network richness) 

% create logical index vectors for plant vs animal species
plants = species_list.groups == "Plants";
animals = species_list.groups ~= "Plants";

% remove consumption of resource types that only plants or animals have
% (can happen from duplication of genus names across taxonomic groups
animal_with_plant_resource_flags = (ismember(expanded_edge_list(:,3),find(~plants))) & (ismember(expanded_edge_list(:,2),find(plant_resource_indx)));
expanded_edge_list(animal_with_plant_resource_flags,:) = [];

% the multiplex food web will be of dimensions (S x S x N)
multiplex_network = zeros(S,S,N);

num_group_species = zeros(N,1);
num_group_animals = zeros(N,1);
num_group_plants = zeros(N,1);
num_group_consumers = zeros(N,1);
num_group_resources = zeros(N,1); 

for i = 1:N

    interaction_group = i;

    group_entries = ismember(expanded_edge_list(:,2), interaction_group);

    group_edge_list = [expanded_edge_list(group_entries,1), expanded_edge_list(group_entries,3)];
    group_web = sparse(group_edge_list(:,1), group_edge_list(:,2), ones(size(group_edge_list,1),1), S, S);

    multiplex_network(:,:,i) = double(logical(full(group_web)));

    % record richnesses of all species and only animals, 
    % plants, consumers, and resources in each subnetwork i
    in_degree_i  = sum(multiplex_network(:,:,i),2);
    out_degree_i = sum(multiplex_network(:,:,i),1)';

    num_group_species(i) = sum(in_degree_i > 0 | out_degree_i > 0); 
    num_group_animals(i) = sum(in_degree_i(animals) > 0 | out_degree_i(animals) > 0);
    num_group_plants(i) = sum(in_degree_i(plants) > 0 | out_degree_i(plants) > 0);
    num_group_consumers(i) = sum(in_degree_i > 0);
    num_group_resources(i) = sum(out_degree_i > 0);

end

% record number of links in each subnetwork 
num_group_links = reshape(sum(sum(multiplex_network,1),2),[N,1]);

num_resources = reshape(sum(multiplex_network,2),[S,N]);
num_consumers = reshape(sum(multiplex_network,1),[S,N]);

species_list.InDegree = num_resources(species_list.indx,:);
species_list.OutDegree = num_consumers(species_list.indx,:);

species_list_degs = species_list;

% analyze the 'traditional' food web where likes are not categorized by
% type (the layers of the multiplex network are 'flattened' into one
% category)
% look at 'flattened_all' which keeps track of when there are double links
flattened_network = sum(multiplex_network,3);
num_species = sum(sum(flattened_network,1) > 0 | (sum(flattened_network,2) > 0)');
num_animals = sum(sum(flattened_network(:,animals),1) > 0 | (sum(flattened_network(animals,:),2) > 0)');
num_plants = sum(sum(flattened_network(:,plants),1) > 0 | (sum(flattened_network(plants,:),2) > 0)');

num_consumers = sum(sum(flattened_network,2) > 0);
num_resources = sum(sum(flattened_network,1) > 0);

richnesses = table(num_group_species, num_group_animals, num_group_plants,...
    num_group_consumers, num_group_resources, 'VariableNames',...
    {'Sx','Ax','Px','Conx','Resx'},'RowNames',interaction_types); 
flattened = table(num_species, num_animals, num_plants, num_consumers, num_resources, ...
    'VariableNames',{'Sx','Ax','Px','Conx','Resx'},'RowNames',{'flattened_all'}); 

% and 'flattened_binary' which keeps track only one link regardless
richnesses = [richnesses; flattened];
linkages = table([num_group_links; sum(logical(flattened_network),'all'); ...
    sum(multiplex_network,'all')],'VariableNames',{'Lx'},...
    'RowNames',{interaction_types{:}, 'flattened_binary','multiplex'});

% count the interactions where consumers  feed on multiple resource types 
% from the same species
multifeeding_type_combos = [];
num_multifeeding_links_on_these_types = [];

% all possible resource combinations
for j = 2:N
    resource_type_combinations = nchoosek(1:N,j);

    for i = 1:size(resource_type_combinations,1)        
        these_resource_types_indx = resource_type_combinations(i,:); 
        these_resource_types_logical = zeros(1,N); 
        these_resource_types_logical(these_resource_types_indx) = 1;

        these_network_layers = multiplex_network(:,:,these_resource_types_indx);

        multifeeding_type_combos = [multifeeding_type_combos; ...
            these_resource_types_logical]; 

        num_multifeeding_links_on_these_types = ...
            [num_multifeeding_links_on_these_types; ...
            sum(sum(these_network_layers,3) == j,'all')]; % this doesn't give the total number of links from this multifeeding but rather the number of double hits

    end
end

multifeeding_links = [multifeeding_type_combos, num_multifeeding_links_on_these_types];
multifeeding_links = splitvars(table(multifeeding_links),'multifeeding_links','NewVariableNames',[interaction_types;"num links"]);

resource_type_combos = [];
num_consumers_eating_these_resource_types = [];
num_plants_supplying_these_resource_types = [];

diet_composition = sum(species_list.InDegree > 0,2);
sp_consumers = diet_composition > 0;
sp_plants = species_list.groups == 'Plants';

% all possible interaction combinations
for j = 1:N
    resource_type_combinations = nchoosek(1:N,j);

    for i = 1:size(resource_type_combinations,1)        
        these_resource_types_indx = resource_type_combinations(i,:); 
        these_resource_types_logical = zeros(1,N); 
        these_resource_types_logical(these_resource_types_indx) = 1;

        resource_type_combos = [resource_type_combos; ...
            these_resource_types_logical]; 

        all_these_resource_types_in = all((species_list.InDegree > 0) == ...
            these_resource_types_logical,2); 

        all_these_resource_types_out = all((species_list.OutDegree > 0) == ...
            these_resource_types_logical,2); 

        num_consumers_eating_these_resource_types = ...
            [num_consumers_eating_these_resource_types; ...
            sum(all_these_resource_types_in(sp_consumers))];

        num_plants_supplying_these_resource_types = ...
            [num_plants_supplying_these_resource_types; ...
            sum(all_these_resource_types_out(sp_plants))];
    end
end

resource_type_usage = table(resource_type_combos, ...
    num_consumers_eating_these_resource_types, num_plants_supplying_these_resource_types, ...
    'VariableNames',{'ResourceCombos','NumConsumers','NumResources'});

resource_type_usage = splitvars(resource_type_usage,'ResourceCombos','NewVariableNames',interaction_types);

end

% function create_diet_breadth_plots(multiplex_network, species_list, data)
% 
% % % CALCULATE TAXONOMIC DIVERSITY OF PLANT RESOURCES
% herbivory_network = multiplex_network(:,:,2:5); 
% 
% [unique_families,list_indx] = unique(species_list.families); 
% unique_families = species_list(list_indx,{'groups','orders','families'});
% 
% for i = 1:height(unique_families)
%     this_family = unique_families.families(i);
%     these_family_members = species_list(strcmp(species_list.families, this_family),:);
%     their_interactions = data(cellfun(@(x) any(ismember(x,these_family_members.indx)),data.ConsumerIndx),:);
%     their_resource_indx = find(sum(sum(herbivory_network(these_family_members.indx,:,:),3),1));
%     their_resources = species_list(arrayfun(@(x) any(ismember(x,their_resource_indx)),species_list.indx),:);
%     their_resource_families = unique(their_resources.families);
%     num_resource_families = numel(their_resource_families);
%     
%     for j = 1:4
%         their_resource_indx_j = find(sum(herbivory_network(these_family_members.indx,:,j),1));
%         their_resources_j = species_list(arrayfun(@(x) any(ismember(x,their_resource_indx_j)),species_list.indx),:);
%         their_resource_families_j = unique(their_resources_j.families);
%         num_resource_families_j(j) = numel(their_resource_families_j);
%     end
%     
%     polyphagy_by_family(i,:) = table(unique_families{i,:},num_resource_families,...
%         num_resource_families_j,'VariableNames',{'family','num_plant_resource_families','num_plant_resource_families_by_type'});
% end
% 
% polyphagy_by_family = splitvars(polyphagy_by_family,'family','NewVariableNames',{'groups','orders','families'});
% 
% % % CREATE IN-DEGREE PLOTS (DIET RICHNESS)
% in_degrees = sum(multiplex_network,2);
% in_degrees = reshape(in_degrees,[3802,5]);
% 
% colors = [ ...
%     169,169,169; ... % gray
%     135,200,182; ... % green
%     226,150,177; ... % pink
%     242, 218, 89; ... % yellow
%     214,169,121; ... % brown
%     ] ./ 256;
% 
% figure
% tiledlayout(3,2)
% set(gcf,'color','w');
% set(gcf,'position',[1,271,490,595])
% 
% nexttile
% i = 1;
% histogram(in_degrees(in_degrees(:,i)>0,i),'FaceColor',colors(i,:),'BinWidth',85)
% set(gca,'FontSize',12);
% xlabel("Num. Animal Resource Species")
% % ylabel("Frequency")
% box off
% size(in_degrees(in_degrees(:,i)>0))
% 
% nexttile
% i = 2;
% histogram(in_degrees(in_degrees(:,3)>0,i),'FaceColor',colors(i,:),'BinWidth',12)
% set(gca,'FontSize',12);
% xlabel("Num. Leaf Resource Species")
% % ylabel("Frequency")
% box off
% size(in_degrees(in_degrees(:,i)>0))
% 
% nexttile
% i = 3;
% histogram(in_degrees(in_degrees(:,2)>0,i),'FaceColor',colors(i,:),'BinWidth',8)
% set(gca,'FontSize',12);
% xlabel("Num. Flower Resource Species")
% % ylabel("Frequency")
% box off
% size(in_degrees(in_degrees(:,i)>0))
% 
% nexttile
% i = 4;
% histogram(in_degrees(in_degrees(:,i)>0,i),'FaceColor',colors(i,:),'BinWidth',8)
% set(gca,'FontSize',12);
% xlabel("Num. Seed Resource Species")
% % ylabel("Frequency")
% box off
% size(in_degrees(in_degrees(:,i)>0))
% 
% nexttile
% i = 5;
% histogram(in_degrees(in_degrees(:,i)>0,i),'FaceColor',colors(i,:),'BinWidth',1)
% set(gca,'FontSize',12);
% xlabel("Num. Wood Resource Species")
% % ylabel("Frequency")
% box off
% size(in_degrees(in_degrees(:,i)>0))
% 
% 
% % load('polyphagy_by_family.mat','polyphagy_by_family')
% polyphagy_by_family = polyphagy_by_family(polyphagy_by_family.groups == "Insects",:);
% 
% nexttile
% h6 = histogram(polyphagy_by_family.num_plant_resource_families ...
%     (polyphagy_by_family.num_plant_resource_families > 0),'BinWidth',1);
% set(gca,'FontSize',12);
% size((polyphagy_by_family.num_plant_resource_families > 0))
% box off
% xlabel("Num. Plant Resource Families")
% % ylabel("Frequency")
% 
% 
% end
% 

function create_degree_correlation_plots(species_list,interaction_types)

in_degrees = table(species_list.InDegree,'VariableNames',{'in_degrees'});
in_degrees = splitvars(in_degrees, {'in_degrees'}, 'NewVariableNames', interaction_types);
in_degrees{:,1:6}(in_degrees{:,1:6}==0) = NaN;

% check correlation coefficient and p-value before plotting detailed
% figures
figure
[R_in,PValue_in] = corrplot(in_degrees,'TestR','on')
title('in degrees')
set(gca,'FontSize',12);
set(gcf,'color','w');

out_degrees = table(species_list.OutDegree,'VariableNames',{'out_degrees'});
out_degrees = splitvars(out_degrees, {'out_degrees'}, 'NewVariableNames', interaction_types);
out_degrees = removevars(out_degrees,"eats animals");
out_degrees{:,1:4}(out_degrees{:,1:4}==0) = NaN;

figure
[R_out,PValue_out] = corrplot(out_degrees,'TestR','on');
title('out degrees')
set(gca,'FontSize',12);
set(gcf,'color','w');

% plot detailed figures (manually using the relationships that are
% significant) and print number of species to terminal
figure
tiledlayout(2,2)
set(gcf,'color','w');

nexttile
scatter(in_degrees.("eats animals"),in_degrees.("eats leaves"),'MarkerEdgeAlpha',0.8)
N11 = sum(~isnan(in_degrees.("eats animals"))& ~isnan(in_degrees.("eats leaves")))

R_in(1,3)
PValue_in(1,3)

xlabel("Animal Resource Richness")
ylabel("Leaf Resource Richness")

h = lsline;
h.Color = [0.8500 0.3250 0.0980];
h.LineWidth = 1;
set(gca,'FontSize',12);

nexttile
scatter(in_degrees.("eats animals"),in_degrees.("eats seeds"),'MarkerEdgeAlpha',0.8)
N12 = sum(~isnan(in_degrees.("eats animals"))& ~isnan(in_degrees.("eats seeds")))
xlabel("Animal Resource Richness")
ylabel("Seed Resource Richness")

R_in(1,4)
PValue_in(1,4)

h = lsline;
h.Color = [0.8500 0.3250 0.0980];
h.LineWidth = 1;
set(gca,'FontSize',12);

nexttile
scatter(in_degrees.("eats leaves"),in_degrees.("eats seeds"),'MarkerEdgeAlpha',0.8)
N21 = sum(~isnan(in_degrees.("eats leaves"))& ~isnan(in_degrees.("eats seeds")))

R_in(3,4)
PValue_in(3,4)

xlabel("Leaf Resource Richness")
ylabel("Seed Resource Richness")

h = lsline;
h.Color = [0.8500 0.3250 0.0980];
h.LineWidth = 1;
set(gca,'FontSize',12);

nexttile
scatter(in_degrees.("eats leaves"),in_degrees.("eats wood"),'MarkerEdgeAlpha',0.8)
N22 = sum(~isnan(in_degrees.("eats leaves"))& ~isnan(in_degrees.("eats wood")))

R_in(3,5)
PValue_in(3,5)

xlabel("Leaf Resource Richness")
ylabel("Wood Resource Richness")

h = lsline;
h.Color = [0.8500 0.3250 0.0980];
h.LineWidth = 1;
set(gca,'FontSize',12);

figure
tiledlayout(2,2)
set(gcf,'color','w');

nexttile
scatter(out_degrees.("eats leaves"),out_degrees.("eats flowers"),'MarkerEdgeAlpha',0.8)
N11 = sum(~isnan(out_degrees.("eats leaves"))& ~isnan(out_degrees.("eats flowers")))

R_out(2,1)
PValue_out(2,1)

xlabel("Leaf Consumer Richness")
ylabel("Flower Consumer Richness")

h = lsline;
h.Color = [0.8500 0.3250 0.0980];
h.LineWidth = 1;
set(gca,'FontSize',12);

nexttile
scatter(out_degrees.("eats leaves"),out_degrees.("eats seeds"),'MarkerEdgeAlpha',0.8)
N12 = sum(~isnan(out_degrees.("eats leaves"))& ~isnan(out_degrees.("eats seeds")))

R_out(2,3)
PValue_out(2,3)

xlabel("Leaf Consumer Richness")
ylabel("Seed Consumer Richness")

h = lsline;
h.Color = [0.8500 0.3250 0.0980];
h.LineWidth = 1;
set(gca,'FontSize',12);

nexttile
scatter(out_degrees.("eats leaves"),out_degrees.("eats wood"),'MarkerEdgeAlpha',0.8)
N21 = sum(~isnan(out_degrees.("eats leaves"))& ~isnan(out_degrees.("eats wood")))

R_out(2,4)
PValue_out(2,4)

xlabel("Leaf Consumer Richness")
ylabel("Wood Consumer Richness")

h = lsline;
h.Color = [0.8500 0.3250 0.0980];
h.LineWidth = 1;
set(gca,'FontSize',12);

nexttile
scatter(out_degrees.("eats flowers"),out_degrees.("eats seeds"),'MarkerEdgeAlpha',0.8)
N22 = sum(~isnan(out_degrees.("eats flowers"))& ~isnan(out_degrees.("eats seeds")))

R_out(1,3)
PValue_out(1,3)

xlabel("Flower Consumer Richness")
ylabel("Seed Consumer Richness")

h = lsline;
h.Color = [0.8500 0.3250 0.0980];
h.LineWidth = 1;
set(gca,'FontSize',12);
end

function check_edge_source(data,species_list,plant_resource_indx)

% full multiplex network can be considered the
% Order Level Network
order_level = data.ConsumerLevel == "Order" ...
    | data.ResourceLevel == "Order";
family_level = data.ConsumerLevel == "Family" ...
    | data.ResourceLevel == "Family";
genus_level = data.ConsumerLevel == "Genus" ...
    | data.ResourceLevel == "Genus"; % acknowledging 285 taxa we could not resolve beyond genus 

species_level = data.ConsumerLevel == "Species" ...
    & data.ResourceLevel == "Species";
all_edge_list = data; 

% build and compare networks at different levels of taxonomic resolution 
% Family Level Network
family_edge_list = data(~order_level,:); 

% Genus Level Network
genus_edge_list = data(~(order_level | family_level),:); 

% Species Level Network
species_edge_list = data(species_level,:); 

% Order ONLY Network
order_only_edge_list = all_edge_list(order_level,:); 

% Family ONLY Network
family_only_edge_list = all_edge_list(family_level & ~order_level,:); 

% Genus ONLY Network
genus_only_edge_list = all_edge_list(genus_level & ~(order_level | family_level),:);

% Build & Analyze Networks
species_list_degs = species_list;

S = height(species_list_degs); 

[species_list_degs, order_richnesses, order_linkages, order_resource_type_usage, ...
    order_multifeeding, order_expanded_edge_list, order_multiplex_network] = analyze_edge_list(all_edge_list, ...
    species_list_degs, plant_resource_indx);
species_list_degs = renamevars(species_list_degs,{'InDegree','OutDegree'},{'InDegree_Order','OutDegree_Order'});

no_links_Order = sum(sum(species_list_degs.InDegree_Order,2) + sum(species_list_degs.OutDegree_Order,2) == 0);

[species_list_degs, family_richnesses, family_linkages, family_resource_type_usage, ...
    family_multifeeding, family_expanded_edge_list, family_multiplex_network] = analyze_edge_list(family_edge_list, ...
    species_list_degs, plant_resource_indx);
species_list_degs = renamevars(species_list_degs,{'InDegree','OutDegree'},{'InDegree_Family','OutDegree_Family'});

no_links_Family = sum(sum(species_list_degs.InDegree_Family,2) + sum(species_list_degs.OutDegree_Family,2) == 0);

[species_list_degs, genus_richnesses, genus_linkages, genus_resource_type_usage, ...
    genus_multifeeding, genus_expanded_edge_list, genus_multiplex_network] = analyze_edge_list(genus_edge_list, ...
    species_list_degs, plant_resource_indx);
species_list_degs = renamevars(species_list_degs,{'InDegree','OutDegree'},{'InDegree_Genus','OutDegree_Genus'});

no_links_Genus = sum(sum(species_list_degs.InDegree_Genus,2) + sum(species_list_degs.OutDegree_Genus,2) == 0);

[species_list_degs, species_richnesses, species_linkages, species_resource_type_usage, ...
    species_multifeeding, species_expanded_edge_list, species_only_multiplex_network] = analyze_edge_list(species_edge_list, ...
    species_list_degs, plant_resource_indx);
species_list_degs = renamevars(species_list_degs,{'InDegree','OutDegree'},{'InDegree_Species','OutDegree_Species'});

no_links_Species = sum(sum(species_list_degs.InDegree_Species,2) + sum(species_list_degs.OutDegree_Species,2) == 0);

[species_list_degs, genus_only_richnesses, genus_only_linkages, genus_only_resource_type_usage, ...
    genus_only_multifeeding, genus_only_expanded_edge_list, genus_only_multiplex_network] = analyze_edge_list(genus_only_edge_list, ...
    species_list_degs, plant_resource_indx);
species_list_degs = renamevars(species_list_degs,{'InDegree','OutDegree'},{'InDegree_GenusOnly','OutDegree_GenusOnly'});

no_links_GenusOnly = sum(sum(species_list_degs.InDegree_GenusOnly,2) + sum(species_list_degs.OutDegree_GenusOnly,2) == 0);

[species_list_degs, family_only_richnesses, family_only_linkages, family_only_resource_type_usage, ...
    family_only_multifeeding, family_only_expanded_edge_list, family_only_multiplex_network] = analyze_edge_list(family_only_edge_list, ...
    species_list_degs, plant_resource_indx);
species_list_degs = renamevars(species_list_degs,{'InDegree','OutDegree'},{'InDegree_FamilyOnly','OutDegree_FamilyOnly'});

no_links_FamilyOnly = sum(sum(species_list_degs.InDegree_FamilyOnly,2) + sum(species_list_degs.OutDegree_FamilyOnly,2) == 0);

[species_list_degs, order_only_richnesses, order_only_linkages, order_only_resource_type_usage, ...
    order_only_multifeeding, order_only_expanded_edge_list, order_only_multiplex_network] = analyze_edge_list(order_only_edge_list, ...
    species_list_degs, plant_resource_indx);
species_list_degs = renamevars(species_list_degs,{'InDegree','OutDegree'},{'InDegree_OrderOnly','OutDegree_OrderOnly'});

family_only_multiplex_network(:,:,5) = zeros(S,S); 

order_only_multiplex_network(:,:,5) = zeros(S,S); 

no_links_OrderOnly = sum(sum(species_list_degs.InDegree_OrderOnly,2) + sum(species_list_degs.OutDegree_OrderOnly,2) == 0);

% fraction of species with links only from order-level records % (ln 478)
(no_links_Family - no_links_Order)/S

% fraction of species with links only from family-level records % (ln 478)
(no_links_Genus - no_links_Family)/S

% fraction of species with links only from genus-level records % (ln 478)
(no_links_Species - no_links_Genus)/S

% Records of feeding on insect orders by any taxon contribute 88% of unique
% carnivory links overall, meaning that they are not otherwise included by 
% records at lower taxonomic levels
sortrows(species_list,'indx','ascend');

generalist_species = sortrows(species_list,'SumInDegree','descend');
generalist_species = generalist_species(1:120,:);

frac_unique_carnivory_links_on_insect_orders = sum(order_only_multiplex_network(:,species_list.groups == "Insects",1) ...
    - family_multiplex_network(:,species_list.groups == "Insects",1) > 0,'all') ...
    / sum(order_multiplex_network(:,:,1),'all') % (ln 343) 

frac_unique_carnivory_links_on_insect_orders_by_generalists = ...
    sum(order_only_multiplex_network(generalist_species.indx,species_list.groups == "Insects",1) ...
    - family_multiplex_network(generalist_species.indx,species_list.groups == "Insects",1) > 0,'all') ...
    / sum(order_multiplex_network(:,:,1),'all') % (ln 339-341)

frac_records_from_generalists_on_insect_orders = sum(ismember(data.Consumer, generalist_species.species_names)...
    & data.ResourceLevel == "Order")/height(data)


frac_unique_carnivory_links_on_insect_orders_by_passeriformes_or_bats = ...
    sum(order_only_multiplex_network(species_list.orders == "Passeriformes" ...
    | species_list.orders == "Chiroptera",species_list.groups == "Insects",1) ...
    - family_multiplex_network(species_list.orders == "Passeriformes" ...
    | species_list.orders == "Chiroptera",species_list.groups == "Insects",1) > 0,'all') ...
    / sum(order_multiplex_network(:,:,1),'all') % (ln 339-341)


frac_records_carnivory_on_insect_orders_by_passeriform_species = ...
    sum((data.ResourceGroup == "Insects" & ...
    data.ResourceLevel == "Order") & ...
    (data.ConsumerLevel == "Species" & ...
    ismember(data.Consumer, ...
    species_list.species_names( ...
    species_list.orders == "Passeriformes")))) / height(data)

tetrapods = species_list(~ismember(species_list.groups,["Insects","Plants","Invertebrates"]),:);

frac_tetrapod_carnivory = sum(order_multiplex_network(tetrapods.indx,tetrapods.indx,1),'all') ...
    / sum(order_multiplex_network(:,:,1),'all')

frac_unique_herbivory_from_orders = sum(order_only_multiplex_network(:,:,2:end) ...
    - family_multiplex_network(:,:,2:5) > 0,'all') ...
    / sum(order_multiplex_network(:,:,2:5),'all') % (ln 343) 

frac_unique_herbivory_from_families = sum(family_only_multiplex_network(:,:,2:5) ...
    - genus_multiplex_network(:,:,2:5) > 0,'all') ...
    / sum(order_multiplex_network(:,:,2:5),'all') % (ln 343) 

frac_unique_herbivory_from_genera = sum(genus_multiplex_network(:,:,2:5),'all') ...
    / sum(order_multiplex_network(:,:,2:5),'all') % (ln 355) 

frac_unique_herbivory_from_genera_by_insects = sum(genus_multiplex_network(species_list.groups == "Insects",:,2:5),'all') ...
    / sum(order_multiplex_network(:,:,2:5),'all') % (ln 356) 

% writetable(species_list_degs,'species_degrees.csv')

end
```
